# A cerebellar cognitive rheostat bidirectionally controls attention

**DOI:** 10.64898/2026.03.22.713532

**Authors:** Shaofei Jiang, Zhaoqi Dong, Zhihong Song, Guochao Yang, Zhouyang Shen, Xuqian Yin, Hao Li, Xinyue Ma, Tiemei Ding, Zhe Zhao, Jun Yang, Huailin Wang, Wei Shen, Huihui Jiang, Liping Chen, Haitao Wu

## Abstract

Attention requires filtering distractors and amplifying signals, processes classically attributed to cortico-thalamic networks. Here, we reveal that the cerebellum operates as a bidirectional “cognitive rheostat” to optimize attentional states. In mice, the anterior and posterior cerebellar vermis exert opposing control over attention. Granule cells in the anterior vermis are functionally suppressed to gate sensorimotor noise via reticular nucleus-driven feedforward inhibition. Conversely, posterior granule cells are recruited by pontine inputs to amplify cognitive signals, a process relying on *Grin1*-mediated NMDA receptor plasticity. Circuit-specific manipulations targeting this push-pull mechanism, or localized pharmacological modulation, successfully rescued attentional deficits in an ADHD mouse model. These findings fundamentally expand the cerebellum’s role beyond motor coordination, identifying a topographic circuit algorithm essential for cognitive control.

## Main Text

Attention depends on two computations that are often discussed together but are rarely mechanistically separated: irrelevant information must be filtered, and task-relevant information must be amplified. Classical accounts have assigned these operations mainly to distributed cortical and thalamic networks, particularly interactions among prefrontal, parietal, sensory, and thalamic association regions (*1-5*). Those networks are essential, but they do not fully explain how attentional state is adjusted rapidly and reliably during ongoing behavior. The cerebellum is well placed to contribute to such control. It contains approximately 80% of the brain’s neurons in total, is linked anatomically to associative cortex, and is implicated in cognitive and affective deficits after injury (*6-9*). Yet the logic by which cerebellar circuits participate in attention remains unresolved, especially at the level of identified lobules, local circuit motifs, and molecular effectors.

Several observations argue that this problem is worth resolving. Human lesion and imaging studies have associated the posterior cerebellar vermis with attentional control and with syndromes in which distractibility and executive dysfunction are prominent (*10-12*). Structural abnormalities of the cerebellum, especially in posterior vermis, are also repeatedly reported in attention-deficit/hyperactivity disorder (ADHD) (*13-17*). At the same time, recent work has expanded the cerebellum’s known repertoire beyond motor coordination, showing that cerebellar granule cells can represent reward expectation, motor planning, and other nonmotor variables (*18-20*). These advances raise an important question: does the cerebellum contribute to attention through a single generic computation across lobules, or do distinct cerebellar territories implement different operations that together set the signal-to-noise ratio (SNR) for attentive behavior?

### Opposing activation of anterior and posterior cerebellar lobules during attention

We addressed this question in mice performing the touchscreen-based 5-choice serial reaction time task (5-CSRTT) (Fig. 1A, fig. S1A to G, Movie S1), a widely used assay of visuospatial attention and response control (*21*). After mice reached criterion performance, brains were collected according to the workflow shown in Fig. 1B, and we first asked whether attentional training engages the cerebellar vermis uniformly. Immunostaining for c-Fos showed selective activation of granule cells in posterior lobules, with the strongest task-associated signal in lobules VI-X (Fig. 1C and D). By contrast, staining for phosphorylated pyruvate dehydrogenase (pPDH), used here as an indicator of suppressed or inactive neurons (*22*), increased predominantly in anterior lobules I-IV (Fig. 1E and F). Thus, the same attentional training that recruited posterior granule cells was accompanied by relative suppression of anterior granule cells. This regional dissociation suggested that the vermis does not simply scale its activity up or down as a whole during attentive behavior.

**Fig. 1.**
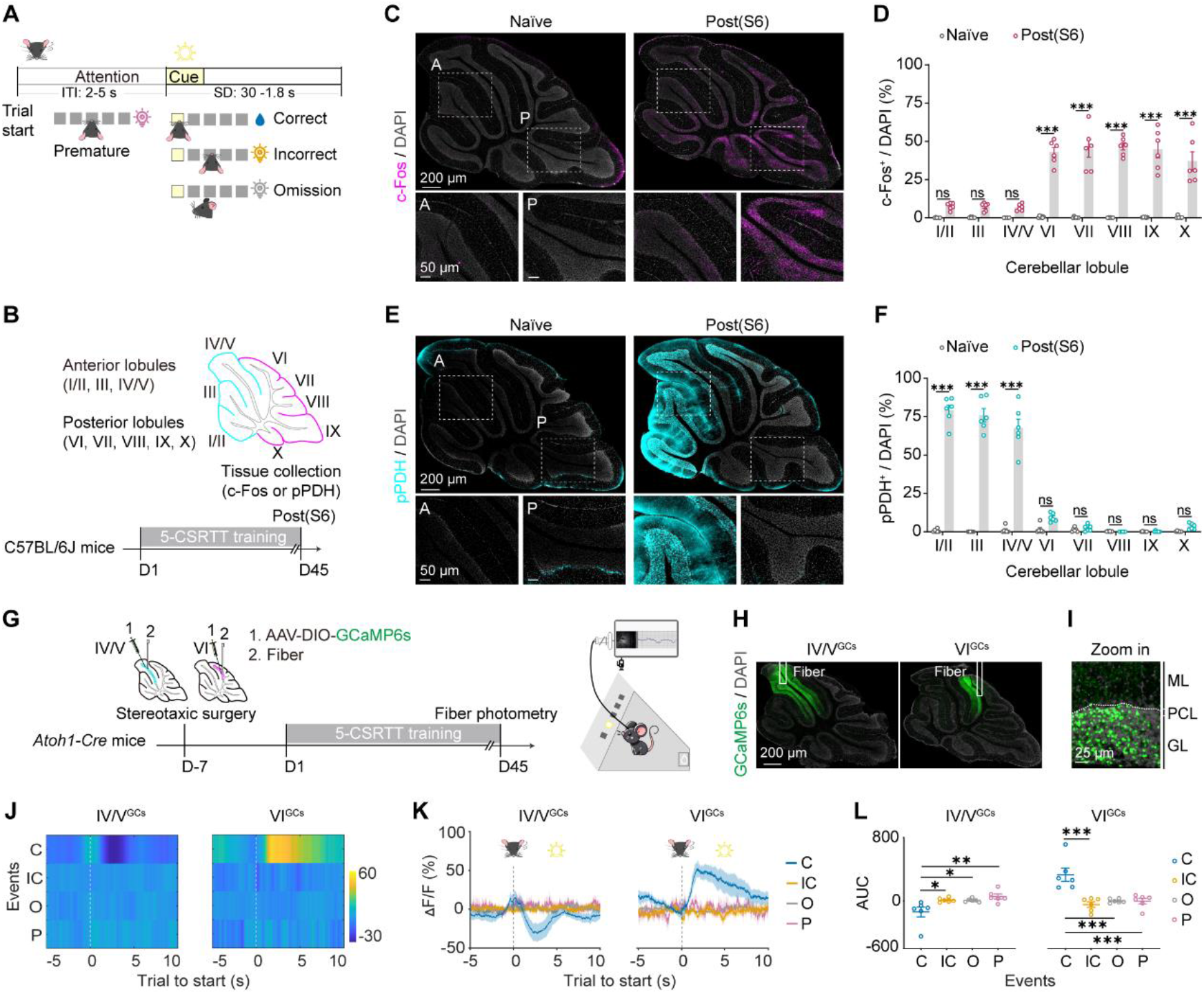
Opposing activity states of anterior and posterior cerebellar granule cells during attentive performance. (A) Schematic of the touchscreen-based 5-choice serial reaction time task (5-CSRTT). Mice initiated a trial, traversed an intertrial interval, and responded to a pseudo-randomly presented visual cue; outcomes were scored as correct, incorrect, omission, or premature. (B) Experimental workflow for activity mapping in naive, pretraining, and trained mice and schematic definition of the anterior and posterior vermis. (C and D) c-Fos staining and quantification show preferential activation of posterior granule cells after training. (E and F) pPDH staining and quantification show preferential suppression of anterior granule cells after training. Scale bars, 200 μm and 50 μm in (C) and (E), as indicated. (G to I) Strategy and validation for in vivo fiber photometry in *Atoh1-Cre* mice expressing GCaMP6s in lobules IV/V or VI. Scale bars, 200 μm in (H) and 25 μm in (I). GL, granular layer; ML, molecular layer; PCL, Purkinje cell layer. (J to L) Heat maps, averaged ΔF/F traces, and area-under-the-curve quantification aligned to trial start show suppression in lobules IV/V and activation in lobule VI during correct trials relative to incorrect, omission, and premature trials. Data are mean ± SEM. n = 6 mice per group for immunostaining in (C to F) and n = 6 mice for photometry in (J to L). Two-way ANOVA with Tukey’s multiple-comparisons test in (D) and (F); one-way ANOVA with Tukey’s multiple-comparisons test in (L). \**P* < 0.05, \*\**P* < 0.01, \*\*\**P* < 0.001.

To resolve the temporal dynamics of these responses, we performed fiber photometry in *Atoh1-Cre* mice expressing GCaMP6s selectively in granule cells (Fig. 1G to I). During correct trials in the 5-CSRTT, calcium signals in lobules IV/V decreased relative to incorrect, omission, and premature trials, whereas lobule VI showed the opposite pattern and increased selectively during correct trials (Fig. 1J to L, Movie S2 and S3). Similar analyses across additional lobules showed little modulation in lobule III but increased signals in lobules VII to IX during correct performance (fig. S2, E to H). These effects were not reproduced by control virus expression (fig. S2, A to D) and were not explained by reward consumption. In naive mice consuming sucrose freely, lobules IV/V and VI showed only small, similar responses (fig. S2, I to L), and reward devaluation in trained mice did not abolish the task-related divergence between the two lobules (fig. S2, M to O). Moreover, the opposing activity pattern was absent before training and emerged progressively as the animals learned the task (fig. S2, P to R). The anterior suppression and posterior recruitment therefore reflected an acquired feature of attentive performance rather than an intrinsic difference in reward processing or movement execution.

### Bidirectional control of attention by granule cells in cerebellar lobules IV/V and VI

We next asked whether these signals merely correlate with behavior or instead contribute causally to performance. We employed chemogenetic manipulation of granule cells in the two lobules to answer this question (Fig. 2A and B). Viral targeting was confined to the intended granule cell populations, and open-field testing showed no detectable change in total distance traveled or time spent in the center across conditions (Fig. 2C to E). During the 5-CSRTT, manipulation of lobules IV/V and VI produced opposite behavioral outcomes. In lobules IV/V, inhibition increased correct responses and correct rate and reduced omissions, whereas activation impaired performance, without changing task velocity (Fig. 2F to I). In lobule VI, the pattern was reversed: inhibition reduced correct responses and correct rate and increased omissions, whereas activation improved performance, again without affecting task velocity (Fig. 2J to M). These findings demonstrate that anterior (lobule IV/V) and posterior (lobule VI) granule cells exert bidirectional, opposing control over attentional performance.

**Fig. 2.**
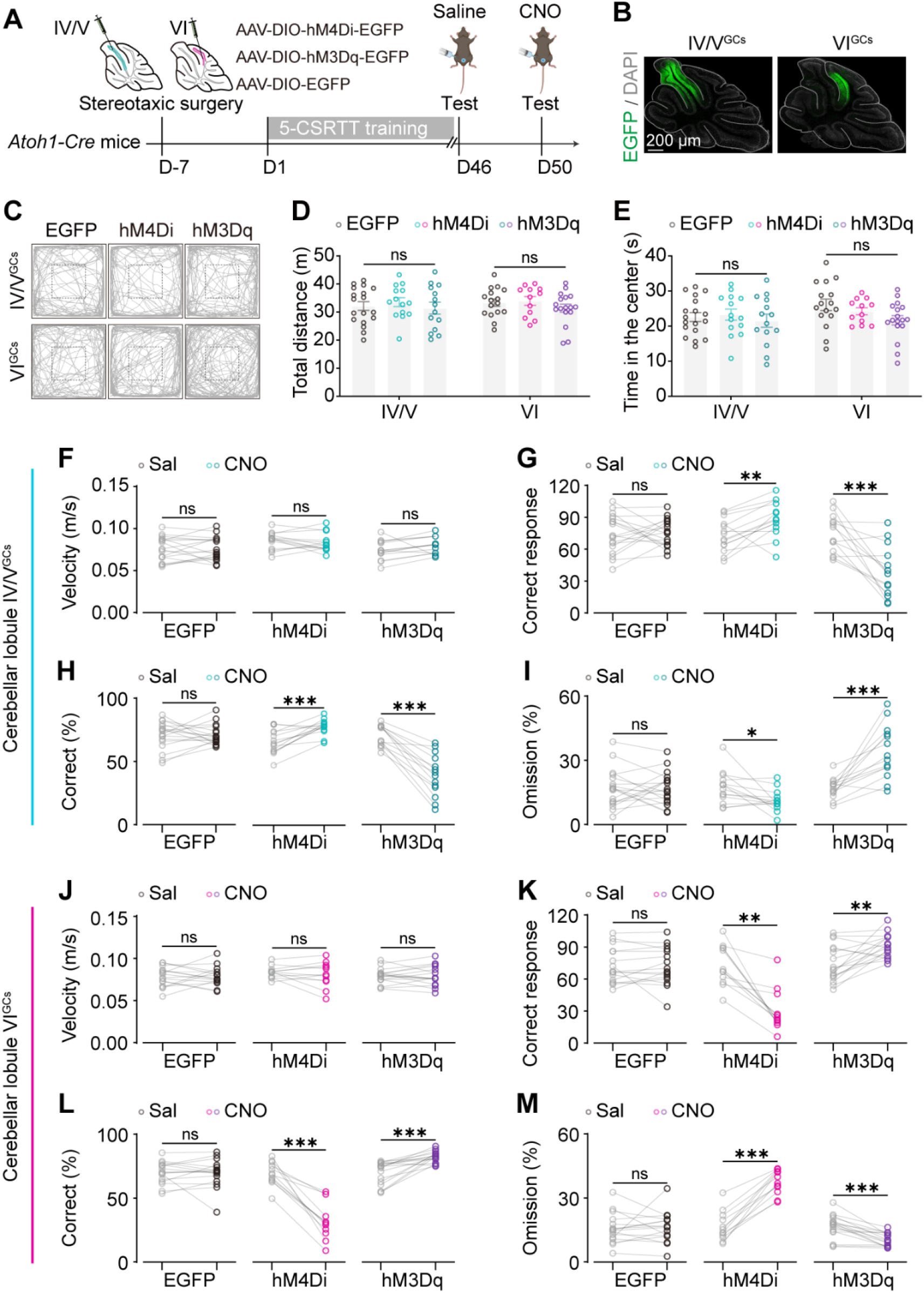
Opposite causal effects of granule cell manipulation in lobules IV/V and VI. (A and B) Chemogenetic strategy and representative viral targeting in *Atoh1-Cre* mice injected with Cre-dependent EGFP, hM4Di, or hM3Dq into lobules IV/V or VI. Scale bar, 200 μm. (C to E) Open-field testing after saline or CNO administration, including representative trajectories (C), total distance traveled (D), and time spent in the center (E). (F to I) Effects of manipulating lobule IV/V granule cells during the 5-CSRTT, shown as task velocity (F), correct responses (G), correct rate (H), and omission rate (I). (J to M) Effects of manipulating lobule VI granule cells during the 5-CSRTT, shown as task velocity (J), correct responses (K), correct rate (L), and omission rate (M). Inhibiting lobule IV/V granule cells improved attention, whereas activating them impaired it; the opposite pattern was observed in lobule VI. Data are mean ± SEM. n = 12 to 18 mice per group. Two-way ANOVA with Tukey’s multiple comparisons test for the open-field measures in (D) and (E); paired t tests for within-animal saline-versus-CNO comparisons in (F to M). \**P* < 0.05, \*\**P* < 0.01, \*\*\**P* < 0.001; ns, not significant.

### Increased expression of *Grin1* in posterior lobule granule cells following the 5-CSRTT

This reversal in granule cell function prompted us to search for molecular differences that might distinguish the two regionally distinct granule cell populations. We therefore performed spatial transcriptomic profiling of the cerebellar vermis in naive mice and mice that had completed 5-CSRTT training (Fig. 3A). Quality-control metrics and marker-gene analysis supported the robustness of the dataset (fig. S3, A to C). Across the dataset, granule cells were the dominant neuronal population and occupied a central position in intercellular communication networks (Fig. 3B to D and fig. S3, D to F). Rank-rank hypergeometric overlap further suggested coordinated transcriptomic changes between lobules IV/V and VI (fig. S3G). However, transcriptional changes after training were not homogeneous across lobules. Granule cells in anterior lobules (ALs) showed enrichment for genes associated with synaptic organization and postsynaptic architecture, whereas granule cells in posterior lobules (PLs) were enriched for genes related to calcium handling, ion transport, and calcium homeostasis (Fig. 3E to G). Among the posterior-enriched candidates, *Grin1* stood out because of its task-associated increase, its central role in NMDA receptor signaling, and its strong functional plausibility for experience-dependent modulation of circuit gain (Fig. 3H).

**Fig. 3.**
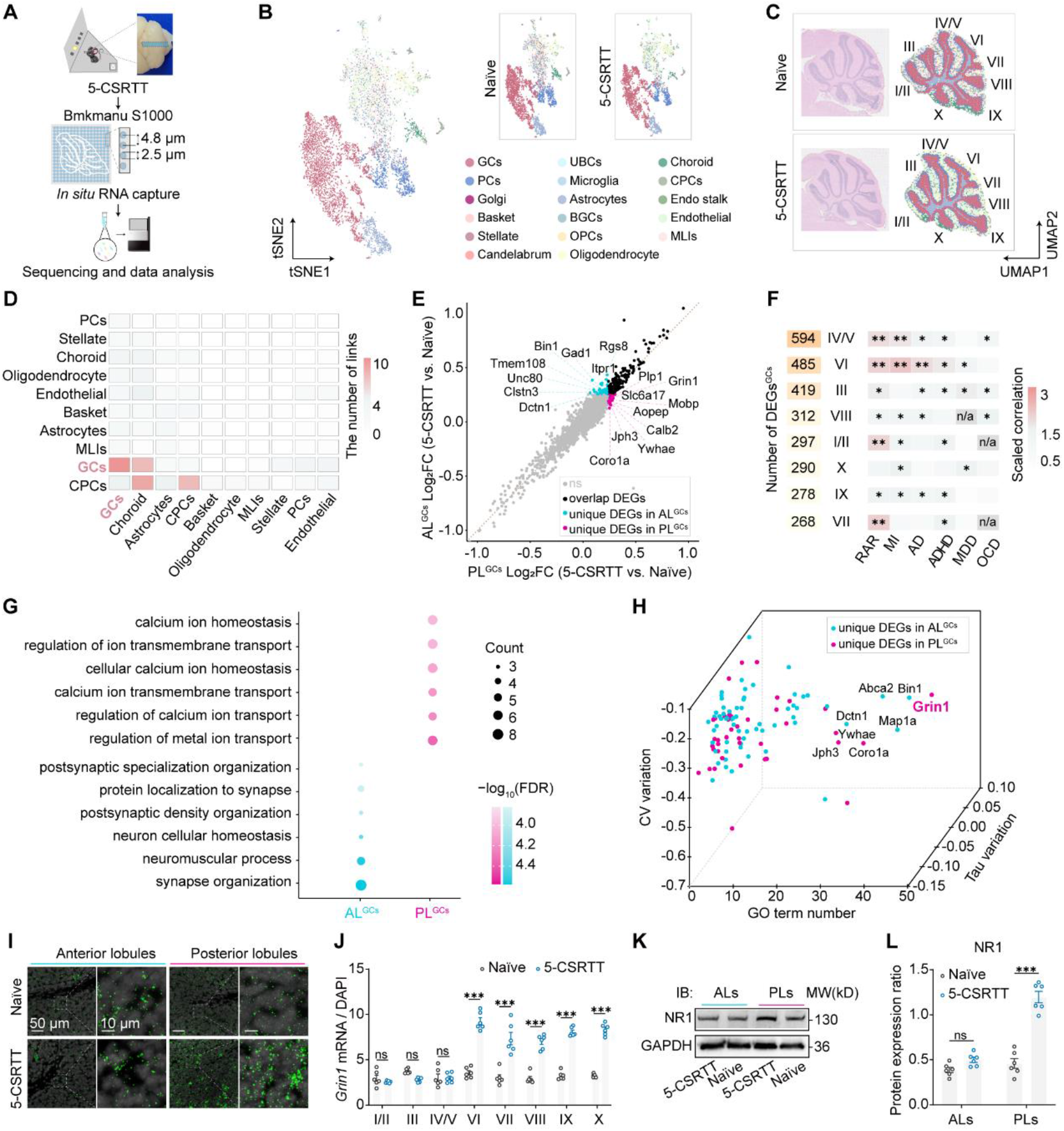
Spatial transcriptomics identifies lobule-specific molecular programs and posterior *Grin1* enrichment. (A) Workflow for spatial transcriptomic profiling of the cerebellar vermis in naive and 5-CSRTT-trained mice. (B and C) Integrated clustering and spatial localization of major cerebellar cell types. (D) Granule cells occupy a central position in inferred intercellular communication networks. (E) Comparison of differentially expressed genes in anterior versus posterior granule cells. (F) Lobule-specific differentially expressed genes and their overlap with neuropsychiatric gene sets. (G) Gene Ontology enrichment of anterior and posterior granule cell programs. (H) Candidate prioritization identifies *Grin1* as a posterior-enriched regulator. (I to L) Validation of posterior *Grin1* enrichment by RNAscope (I and J) and immunoblotting of NR1 protein (K and L). Scale bars, 50 μm (low view) and 10 μm (high view) in (I), as indicated. Data are mean ± SEM. n = 6 mice for (J) and (L). Two -way ANOVA with Sidak’s multiple-comparisons test in (J); two-way ANOVA with Tukey’s multiple-comparisons test in (L). \**P* < 0.05, \*\**P* < 0.01, \*\*\**P* < 0.001; ns, not significant.

We validated the transcriptomic result by RNAscope analysis and immunoblotting. Both *Grin1* mRNA and the encoded NR1 protein increased selectively in posterior lobules after 5-CSRTT training, whereas the anterior vermis showed no comparable change (Fig. 3I to L). We then tested whether local NMDAR signaling in the two lobules is behaviorally required using site-specific infusion of the noncompetitive antagonist MK-801 (*23, 24*). Open-field titration showed that 20 nmol did not measurably change total distance traveled, whereas 40 nmol reduced time spent in the center, so 20 nmol was used for task experiments (Fig. 4A to C). Cannula placement and the testing schedule are shown in Fig. 4D and E. In lobule VI, both acute and repeated MK-801 treatment reduced correct responses and correct rate and increased omissions without changing task velocity (Fig. 4F to I). In contrast, acute antagonism in lobules IV/V had little effect, whereas repeated antagonism improved performance, increasing correct responses and correct rate and decreasing omissions without affecting velocity (Fig. 4J to M). These results identify posterior-lobule VI NMDAR signaling, and particularly *Grin1*-linked signaling, as a key molecular requirement for the pro-attentional function of this region.

**Fig. 4.**
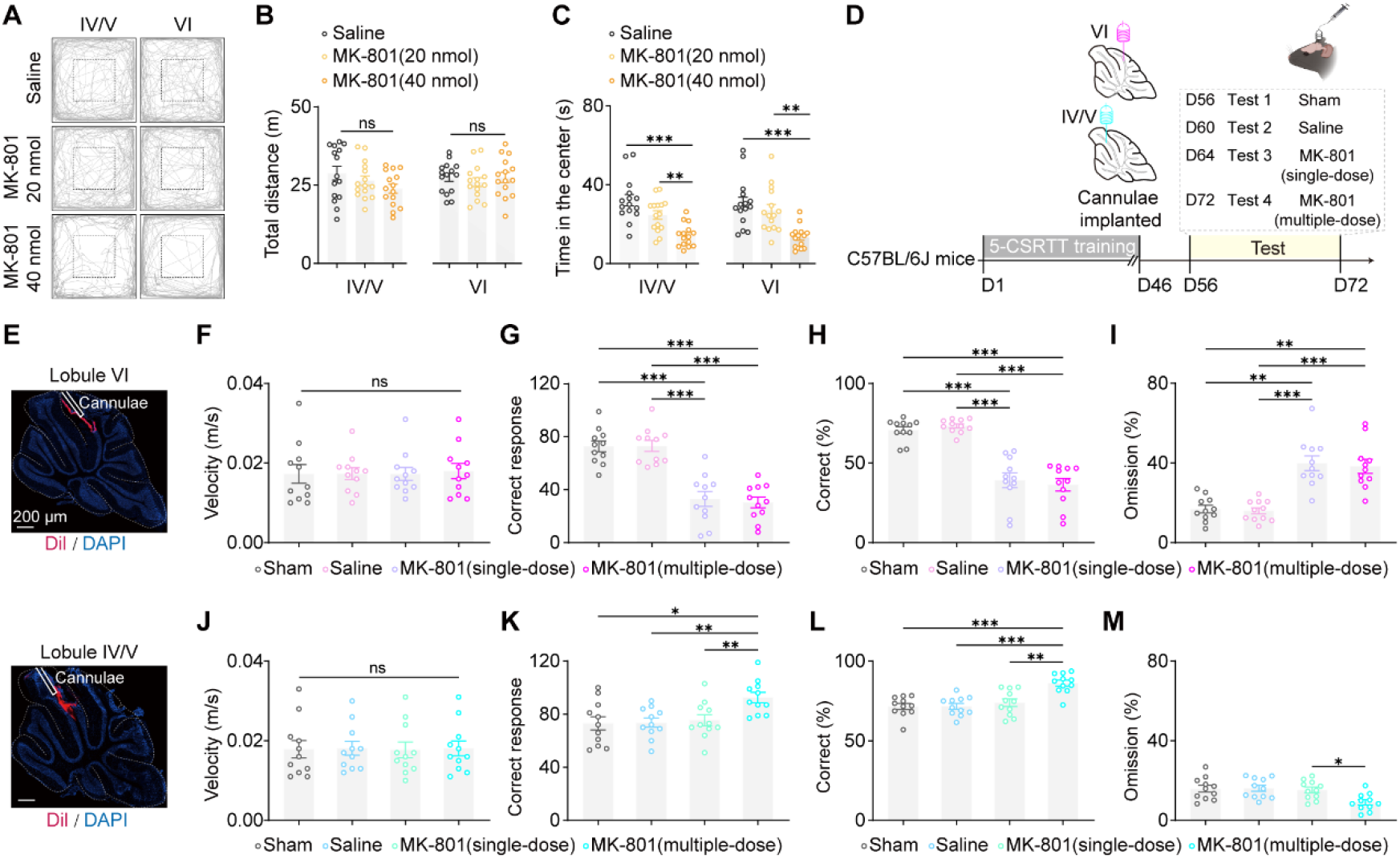
Lobule-specific NMDA receptor signaling contributes to attentional performance. (A to C) Open-field dose titration for local MK-801 infusion into lobules IV/V or VI, including representative locomotor trajectories (A), total distance traveled (B), and time spent in the center (C). The 20-nmol dose did not measurably alter locomotion, whereas the 40-nmol dose reduced center time. (D) Experimental design for sequential sham, saline, single-dose, and multiple-dose MK-801 testing during the 5-CSRTT. (E) Representative sagittal sections showing cannula placement in lobule VI (upper) and lobules IV/V (lower). Scale bar, 200 μm. (F to I) Effects of MK-801 infusion into lobule VI on task velocity (F), correct responses (G), correct rate (H), and omission rate (I). (J to M) Effects of MK-801 infusion into lobules IV/V on task velocity (J), correct responses (K), correct rate (L), and omission rate (M). Data are mean ± SEM. n = 15 mice per group for dose titration and n = 11 mice per group for the repeated-measures 5-CSRTT analyses. One-way ANOVA with Tukey’s multiple-comparisons test in (B) and (C); repeated-measures ANOVA with Tukey’s multiple-comparisons test in (F to M). \**P* < 0.05, \*\**P* < 0.01, \*\*\**P* < 0.001; ns, not significant.

### Granule cells in lobule IV/V and lobule VI receive distinct projections and exhibit specific connectivity patterns

The opposing causal roles of lobules IV/V and VI raised a second question: are these territories embedded in different afferent architectures? We addressed this with monosynaptic rabies tracing from granule cells in each lobule (fig. S4, A to E) (*25*). Lobule VI received prominent inputs from pontine nuclei and the reticulotegmental nucleus, consistent with relay of cortical information to posterior cerebellum. In contrast, lobules IV/V received a broader set of inputs from reticular and sensorimotor-related nuclei, including the reticular nucleus, gigantocellular reticular nucleus, vestibular nuclei, and cuneate nucleus. Anterograde tracing confirmed this topography: pontine axons preferentially innervated posterior lobule VI, whereas reticular and vestibular inputs were concentrated in anterior lobules IV/V (fig. S4, F and G). The anatomical divergence suggested that the reversal in granule cell function might be implemented by distinct upstream pathways rather than by a common input interpreted differently by local circuitry.

### Distinct cerebellar afferent pathways causally mediate attention control

To determine which of these afferents are engaged during attention, we combined retrograde labeling from cerebellar cortex with c-Fos mapping in upstream nuclei after the 5-CSRTT (fig. S5, A to D). Projection neurons in pontine nuclei that targeted lobule VI, and projection neurons in reticular nucleus that targeted lobules IV/V, were robustly recruited during task performance. By contrast, lateral vestibular neurons projecting to lobules IV/V showed little or no task-dependent recruitment. We then selectively manipulated each pathway with an intersectional chemogenetic strategy (Fig. 5A and B). Inhibition of the pontine-to-lobule VI pathway impaired attentional performance, whereas activation enhanced it (Fig. 5C to E and L to N). Manipulating the lateral vestibular-to-lobule IV/V pathway had no measurable effect (Fig. 5F to H and O to Q). By contrast, inhibition of the reticular-to-lobule IV/V pathway worsened attention, whereas activation improved it (Fig. 5I to K and R to T). None of these manipulations altered task velocity (fig. S5E and F).

**Fig. 5.**
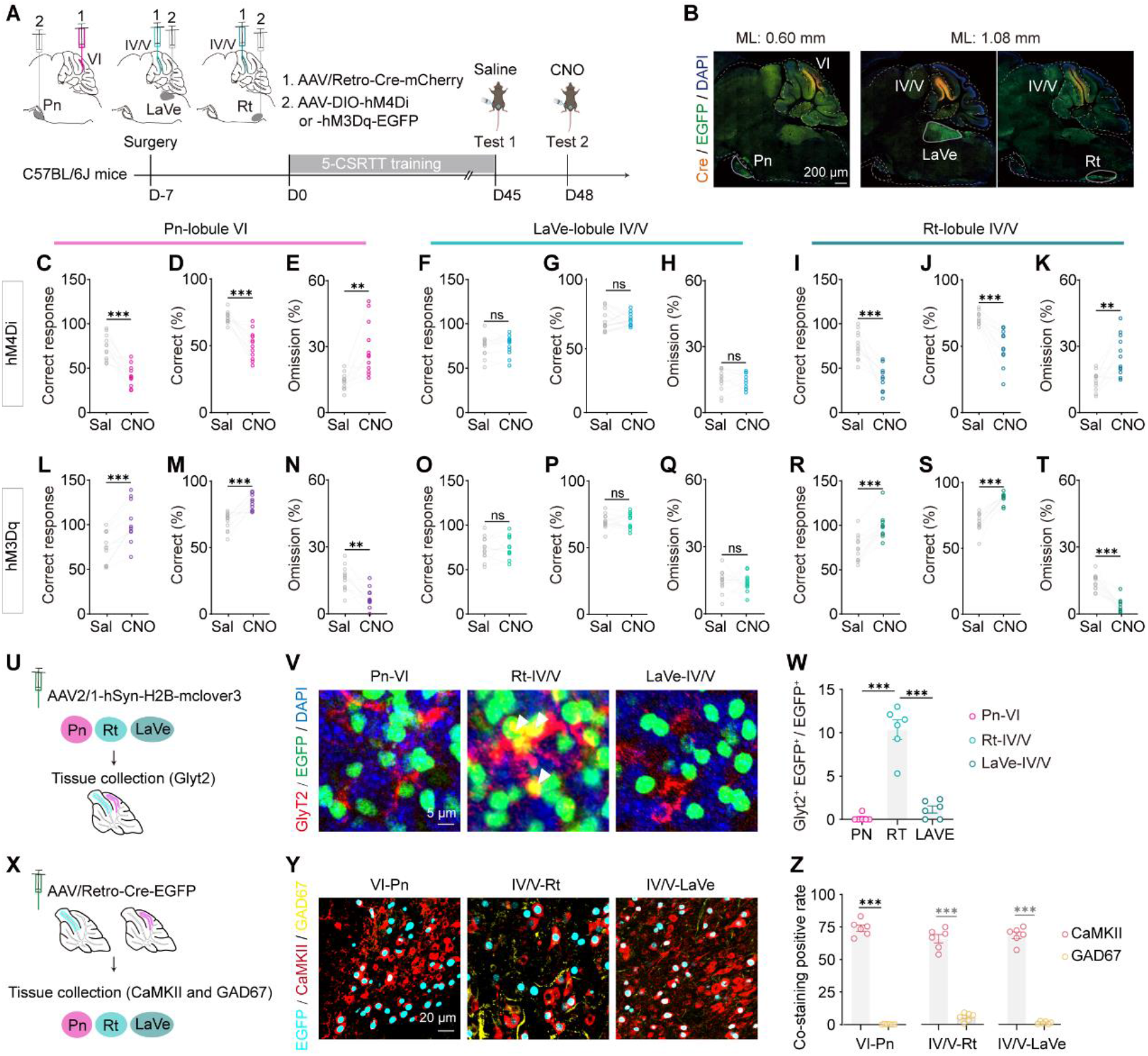
Distinct cerebellar afferent pathways implement anterior suppression and posterior amplification. (A and B) Intersectional strategy and representative expression for pathway-specific chemogenetic manipulation of pontine nuclei (Pn), lateral vestibular nucleus (LaVe), and reticular nucleus (Rt) inputs to the cerebellum. Scale bar, 200 μm. (C to E and L to N) Manipulation of the Pn-to-lobule VI pathway. Inhibition impaired attention (C to E), whereas activation improved it (L to N). (F to H and O to Q) Manipulation of the LaVe-to-lobule IV/V pathway had no measurable effect. (I to K and R to T) Manipulation of the Rt-to-lobule IV/V pathway bidirectionally regulated performance, with inhibition impairing and activation improving attention. (U) Experimental design for assessing synaptic organization. (V and W) Representative confocal images and quantification of afferent terminals in relation to GlyT2-positive Golgi cells. Scale bars, 5 μm in (V). (X to Z) Strategy, representative images, and quantification used to define the neurotransmitter identity of projection neurons. Scale bars, 20 μm in (Y). Data are mean ± SEM. n = 12 mice for behavioral experiments and n = 6 mice for histological analyses. Paired t tests in (C to T); one-way ANOVA with Tukey’s multiple-comparisons test in (W); unpaired t test in (Z). \**P* < 0.05, \*\**P* < 0.01, \*\*\**P* < 0.001; ns, not significant.

At first sight, the reticular result appears paradoxical. Granule cells in lobules IV/V are suppressed during correct trials (Fig. 1J to L), yet activation of their reticular input improves performance (Fig. 5R to T). A plausible resolution is that reticular drive does not primarily excite anterior granule cells directly, but instead recruits local inhibition that reduces net granule cell output. Several observations support this interpretation. Reticular terminals in lobules IV/V showed substantially greater structural apposition to GlyT2^+^ Golgi cells than did pontine or vestibular terminals in their target lobules (Fig. 5U to W). Retrograde profiling further indicated that the relevant reticular projection neurons are predominantly excitatory, not GABAergic, implying that the inhibitory effect is likely mediated locally (Fig. 5X to Z). Together, these findings support a feedforward model in which excitatory reticular afferents engage Golgi cell inhibition in anterior vermis, thereby suppressing granule cell activity and limiting sensorimotor interference during correct performance. This interpretation is consistent with general principles by which feedforward inhibition sharpens timing, normalizes gain, and suppresses competing activity (*26-28*).

This arrangement differs sharply from the posterior pathway. Pontine inputs to lobule VI showed much weaker association with Golgi cells (Fig. 5U to W) and instead matched the task-dependent recruitment and positive causal contribution of posterior granule cells (Fig. 1J to L; Fig. 5C to E and L to N). The posterior vermis therefore appears positioned to amplify task-relevant information relayed from cortex through pontine nuclei, whereas the anterior vermis appears organized to suppress behaviorally competing input streams through a local inhibitory motif. In this framework, attention is supported by a push-pull cerebellar architecture: anterior lobules reduce noise, and posterior lobules enhance signal. Retrograde tracing further showed that both pontine and reticular nuclei receive convergent input from medial prefrontal, anterior cingulate, insular, parietal association, and somatosensory cortical territories, placing these cerebellar pathways within a broader network already implicated in attentional control (fig. S6).

### Circuit and molecular interventions rescue attention in an ADHD mouse model

We then asked whether this architecture has translational relevance in a disease model with impaired attention. Dopamine transporter heterozygous knockout mice (*DAT-HET*), a widely used model related to ADHD-like phenotypes (*29-31*), showed hyperactivity in the open field (Fig. 6A to C) and marked deficits in the 5-CSRTT, including fewer correct responses, lower correct rate, and more omissions and premature responses (Fig. 6D to G). We next tested whether activating either of the two cerebellar pathways identified above could improve performance in these mice. Using the intersectional chemogenetic strategy shown in Fig. 6H, stimulation of the reticular- to-lobule IV/V projection restored correct responses and reduced omissions (Fig. 6I to K), and activation of the pontine-to-lobule VI projection produced a comparable rescue (Fig. 6L to N). Because posterior *Grin1*/NMDAR signaling was selectively linked to trained attentive performance, we also tested a molecular intervention. Local NMDA infusion into lobule VI, according to the scheme in Fig. 6O, significantly improved task performance in *DAT-HET* mice (Fig. 6P to R). The rescue by both circuit-level and molecular manipulations indicates that these cerebellar mechanisms are not merely correlated with normal performance but are sufficient to alleviate attention deficits in a disease-relevant background.

**Fig. 6.**
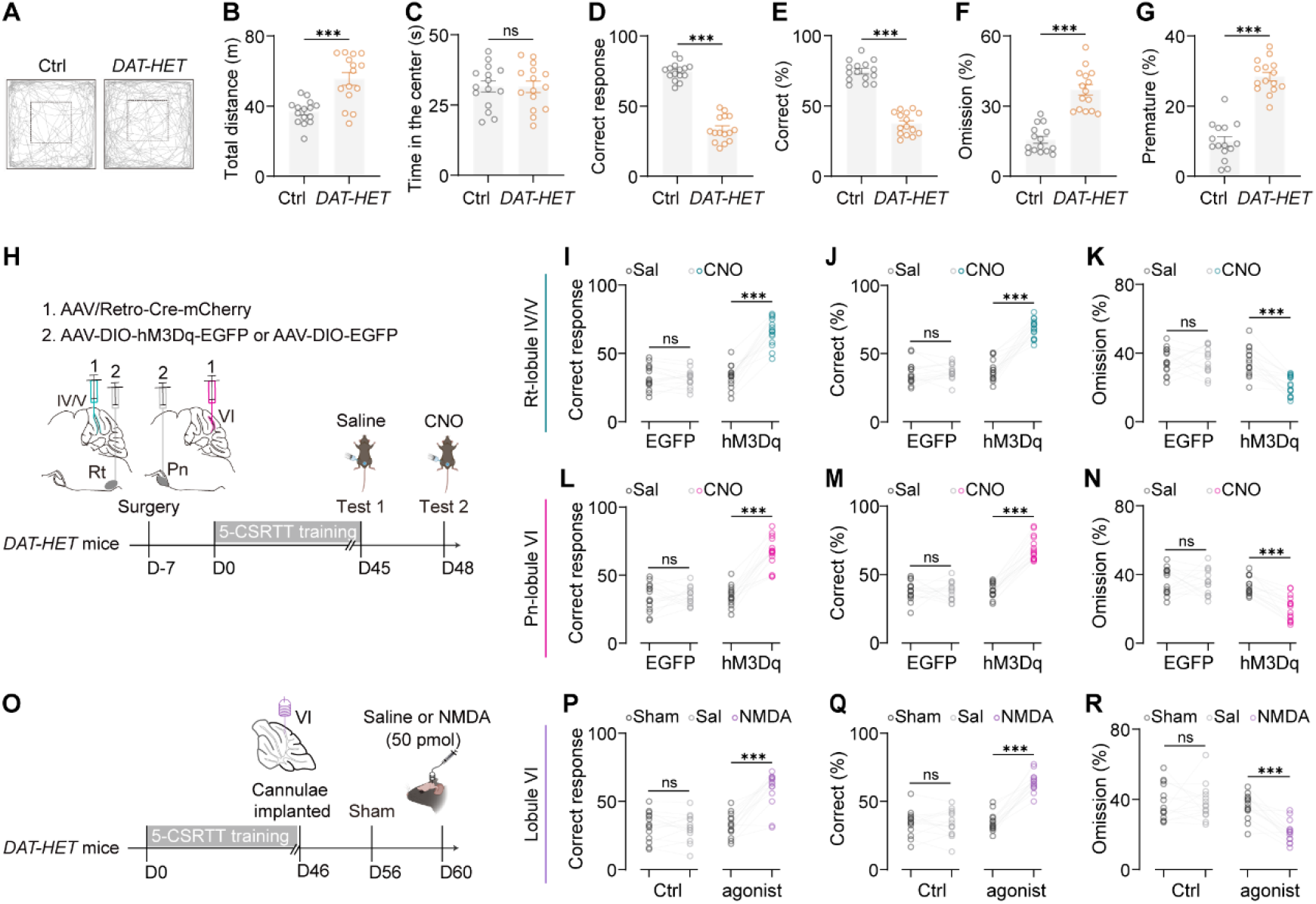
Circuit-level and molecular interventions rescue attention deficits in *DAT-HET* mice. (A to C) Open-field phenotype of control and *DAT-HET* mice, shown as representative trajectories (A), total distance traveled (B), and time spent in the center (C). (D to G) 5-CSRTT phenotype of *DAT-HET* mice, shown as correct responses (D), correct rate (E), omission rate (F), and premature rate (G). (H) Intersectional chemogenetic strategy for pathway-specific rescue. (I to K) Activating the Rt-to-lobule IV/V pathway improved performance in *DAT-HET* mice. (L to N) Activating the Pn-to-lobule VI pathway improved performance. (O) Experimental design for local NMDA infusion into lobule VI. (P to R) Local NMDA improved performance in *DAT-HET* mice. Data are mean ± SEM. n = 15 mice per group for the baseline phenotype, n = 15 mice per group for the chemogenetic rescue experiments, and n = 14 mice per group for NMDA rescue. Unpaired t tests in (B to G); paired t tests in (I to N) and (P to R). \**P* < 0.05, \*\**P* < 0.01, \*\*\**P* < 0.001; ns, not significant.

## Discussion

Attention selectively processes relevant information while filtering distractors (*1, 32*). The prevailing corticocentric view posits that top-down attention is orchestrated primarily by prefrontal-thalamic loops (*2, 4, 33-45*). While critical, this framework struggles to explain rapid gain control and the profound attentional deficits seen in cerebellar pathologies (*8, 11*). Our study identifies the cerebellar vermis as a topographically organized regulator of attention. We demonstrate a dual-process “Push-Pull” mechanism: the anterior lobule (IV/V) is suppressed to gate sensorimotor noise, while the posterior lobule (VI) is recruited to amplify cognitive signals. This cognitive rheostat optimizes the brain’s SNR, providing a physiological basis for the cerebellar cognitive affective syndrome (*8, 11*) and expanding the cerebellum’s role into cognitive control.

A major conceptual advance is resolving a functional paradox in the anterior cerebellum. Granule cells in Lobule IV/V are suppressed during correct attentional performance, despite receiving dense excitatory mossy fibers from the reticular nucleus—a Reticular Activating System (RAS) hub. This silencing occurs via a specialized feedforward inhibitory motif: glutamatergic reticular nucleus afferents preferentially recruit local GlyT2^+^ Golgi cells, which subsequently inhibit granule cells. Because the anterior cerebellum receives dense proprioceptive and motor-planning inputs (*46, 47*), this task-irrelevant sensorimotor stream acts as competing background noise. By utilizing reticular nucleus-driven feedforward inhibition, the brain gates this throughput, enforcing temporal fidelity (*26-28, 48*). Furthermore, while both the reticular nucleus and lateral vestibular nucleus provide sensorimotor inputs, only the reticular nucleus pathway is functionally critical for this gating. This targeted suppression of reticular nucleus-mediated sensorimotor arousal prevents motor overflow and refines our understanding of the RAS. Consistent with human imaging (*49*), this mechanism enables “focused arousal”—energizing the cortex while dampening cerebellar noise.

In contrast, the posterior vermis (Lobule VI) acts as a cognitive amplifier. Its granule cells are robustly activated during successful performance and are innervated by the pontine nuclei, which relay prefrontal, parietal, and cingulate inputs. This suggests a hierarchical organization: the anterior cerebellum processes sensorimotor feedback, while the posterior integrates into large-scale cognitive networks (*50, 51*). This posterior activation likely enhances representations of task-relevant stimuli. By integrating top-down cortical signals, it may calculate attentional error signals, aligning with the Universal Cerebellar Transform theory (*52*). This cerebro-cerebellar communication (Pontine → Lobule VI) maintains the attentional set and aligns with primate data showing cerebellar dentate nucleus neurons encode visual attention (*53*). Ultimately, the cerebellum actively boosts signals (posteriorly) while filtering noise (anteriorly).

Spatial transcriptomics uncovered the molecular logic behind these functional divergences. Anterior granule cells are enriched for synaptic organization genes, whereas posterior granule cells exhibit calcium signaling profiles. Crucially, *Grin1* (encoding the NMDA receptor subunit NR1) is specifically upregulated in the posterior vermis following training. Because NMDA receptors gate plasticity, this cognitive amplifier likely relies on NMDAR-mediated plasticity to strengthen cortico-cerebellar synapses. *Grin1* is also a high-confidence risk gene for ASD and schizophrenia (*54*), both marked by attentional dysfunction. Pharmacological NMDAR inhibition in Lobule VI mimics these deficits, while activation rescues them, directly linking cerebellar molecular pathways to clinical phenotypes and explaining region-dependent disruptions caused by global NMDAR antagonists (*23, 24*).

More importantly, these findings offer a circuit-level framework for ADHD pathophysiology. ADHD patients show reduced vermal volume and altered connectivity (*10, 12-14, 17*). We propose ADHD symptoms reflect a dysregulated push-pull system: failed anterior gating permits sensorimotor noise, driving hyperactivity/impulsivity, while a hypoactive posterior amplifier weakens stimulus representation, causing distractibility. Supporting this, boosting either pathway rescued attentional impairments in the *DAT-HET*, ADHD mouse model (*29-31*). Unlike global treatments like methylphenidate, our work highlights precise interventions, such as targeted neuromodulation (TMS/tDCS) or posterior-specific NMDAR positive allosteric modulators, to treat the “Dysmetria of Thought” (*55*).

In conclusion, our results extend the Universal Cerebellar Transform theory (*52, 55*). Despite conserved anatomy, functional outcomes are dictated by distinct connectivity and molecular identities. Analogous to motor systems canceling self-generated sensations, anterior suppression acts as “attentional attenuation” of predictable noise, while posterior activation enhances signal gain to optimize SNR. Limitations include our exclusive focus on the vermis due to its ADHD implications (*10, 12-14, 17*); lateral hemispheres also likely contribute via prefrontal loops (*55, 56*). Future studies could map these lateral contributions and dissect specific cortical inputs to the reticular nucleus and pontine nuclei. Finally, while the *DAT-HET* mouse is a robust model (*29-31*), capturing human attention’s full complexity requires comparative neuroimaging and clinical trials. By defining this cognitive rheostat, our study offers precise neuropsychiatric targets and solidifies the cerebellum as a critical node in higher-order cognition.

## Acknowledgements

We thank all members of the Wu and Dong laboratories for reading the manuscript.

## Funding

This work was supported by the National Science Fund for Distinguished Young Scholars (32325025 to H.W.), the National Key Research and Development Program of China (2021YFA1101801 to H.W.), the Ministry of Science and Technology Innovation (STI) 2030-Major Projects (2021ZD0202500 to H.W. and 2025ZD0218000 to Z.D.), the National Natural Science Foundation of China (32171148 to H.W. and 32300817 to Z.D.), and the Innovative Group Cultivation Project for Basic Medicine (CX25XT03 to Z.D.).

### Author contributions

H.W. and Z.D. conceived and supervised the study. S.J. and Z.D. performed behavioral assays, fiber photometry, circuit mapping, and chemogenetic experiments. S.J. and Z.S. performed spatial transcriptomic analyses. S.J., G.Y., and H.L. performed immunoblotting, immunohistochemistry, and RNAscope experiments. Z.S., X.Y., X.M., T.D., Z.Z., J.Y., H.W., W.S., H.J., and L.C. provided technical support. S.J., Z.D., and H.W. wrote the manuscript with input from all authors.

## Data and materials availability

Spatial transcriptomic sequencing data have been deposited in the Genome Sequence Archive at the Beijing Institute of Genomics under accession CRA036905. Analysis code is available at https://github.com/songzh523/Spatial_transcriptomics_analysis_of_cerebellar-vermis_after_5-CSRTT. Additional information is available from the corresponding authors upon reasonable request.

