## Supplementary Materials for "A cerebellar cognitive rheostat bidirectionally controls attention"

**The PDF file includes:**

Materials and Methods

Figs. S1 to S6

Tables S1 to S2

**Other Supplementary Materials for this manuscript include the following:**

Movies S1 to S3

Data S1

**Materials and Methods**

**Animal models**

The C57BL/6J mice were obtained from SPF Biotechnology Co., Ltd. (Beijing, China). *Atoh1-Cre* mice were provided by the Jackson Laboratory (stock number 011104) for expression in GCs as previously described (*19, 20*). The *DAT-HET* mice were obtained from Shanghai Model Organisms Center, Inc. Mouse strains and genotyping primer pairs used in this study were all listed in table S1.

Male and female mice (aged 7-8 weeks) were randomly selected for the experiments, with no gender differences observed in the findings. The mice were group-housed (four to five per cage) under a 12-hour light/dark cycle with food and water available *ad libitum* until the start of 5-CSRTT training, at which point they were placed on a water-restricted regimen as described below. All procedures adhered to animal care and biosafety guidelines approved by the Institutional Animal Care and Use Committee of the Beijing Institute of Basic Medical Sciences (#SYXK2019-0004).

**Behavioral analysis**

*The Five-Choice Serial Reaction Time Task (5-CSRTT)*

Mice were trained using the 5-CSRTT procedure according to the published protocol (*57*). Sessions were conducted in standard chambers designed for the 5-CSRTT (Lafayette Instrument Company), within sound-attenuating, ventilated enclosures. The session programs were controlled, and data were collected through a PC-interface setup with the Lafayette Instruments control unit (Version 21.02.26.0, Lafayette Instrument Company), including ABET II Touch and Whisker software (Cambridge University Technical Services Ltd).

Prior to 5-CSRTT training, the mice’s daily water access was gradually reduced from 4 hours to 0.5 hours, with a reduction of 0.5 hours per day. After 8 days of water restriction adaptation, the mice were placed on a water-restricted regimen of 1 mL/day following behavioral sessions, and the 5-CSRTT training began. Mice were monitored daily and weighed to maintain their body weight at approximately 90% of their free-drinking weight to ensure overall health and adequate motivation for task learning. Training sessions were limited to 100 trials or 30 minutes, whichever came first, with training occurring 6 days per week.

Initially, mice were trained to associate the touch screen with a reward (2% sucrose solution). Subsequently, they were trained to respond to one of five touchscreens within a decreasing time limit (from 30 s to 5 s) to earn a 20 μL reward. Incorrect response (e.g., responding to the wrong screen), omission response (failure to respond within the allotted time), or premature response (responding before the stimulus appeared) were followed by a 2-second white houselight illumination and a 5-second timeout. Mice progressed through the stages of training sequentially, advancing upon meeting specific performance criteria (detail see Figure S1A). Animals were excluded from the analysis if their baseline performance prior to testing failed to meet the predefined inclusion criteria. During the test phase, mice could complete trials without a time limit within the 30-minute session. Attention was assessed based on several parameters: correct response, incorrect response, omission, percentage of correct response ([correct / (correct + incorrect + omission)] × 100), percentage of omission ([omission / (correct + incorrect + omission)] × 100), accuracy ([correct / (correct + incorrect)] × 100), and percentage of premature response ([premature / (correct + incorrect + omission + premature)] × 100).

*The Open-Field Test (OFT)*

The OFT can assess motor ability (total distance traveled and velocity) and anxiety levels (time spent in the center) in mice (*58*). Mice were individually placed in an open-field apparatus (custom-made white acrylic box, 50 × 50 × 40 cm) allowing free exploration, and their behavior was recorded for 5 minutes using behavior tracking software (ANY-maze, Stoelting).

*The Novel Object Recognition Test (NORT)*

The Novel Object Recognition Test (NORT) assesses the cognitive memory ability of mice by comparing their exploration time of familiar versus novel objects, as rodents naturally tend to spend more time exploring new objects than familiar ones (*59, 60*). The test is divided into three stages, with mice allowed to explore freely for 5 minutes in each stage. In the first stage, there are no objects in the custom-made acrylic box (80 × 40 × 40 cm). In the second stage, two identical rectangular prisms (5 × 5 × 10 cm) are placed diagonally in the box. In the third stage, one of the rectangular prisms is replaced with a cone (5 cm in diameter) in the left corner of the box. The exploration time of the animals towards different objects is recorded using ANY-maze software, and the object discrimination index (ODI) is calculated in the third stage. The formula for the ODI is: ODI = new object exploration time / (new object exploration time + old object exploration time).

**Immunostaining**

Evaluation of c-Fos and pPDH expression in cerebellar GCs was conducted via immunofluorescence staining (IF). Mice were sacrificed either 1.5 h (*61*) or 15 min (*22*) after the 5-CSRTT, respectively. Mice were anesthetized with 1% pentobarbital (50 mg/kg; P3761, Sigma) and perfused first with saline, followed by pre-cooled 4% paraformaldehyde (PFA). The brains were post-fixed in 4% PFA at 4°C for 24 h, then transferred to 15% and 30% sucrose solutions at 4°C for 24 h each. Sagittal sections of the cerebellar vermis were cut at 40 μm thickness using a cryotome (Thermo Fisher Scientific).

For immunofluorescence, tissue sections were incubated in a blocking buffer consisting of 0.3% Triton X-100 and 3% bovine serum albumin in phosphate-buffered saline (PBS) for 1.5 hours at room temperature (23-24°C). Primary antibodies, diluted in the blocking buffer, were applied and the slides were incubated overnight at 4°C. The following primary antibodies were used: rabbit anti-c-Fos (ab190289, Abcam), rabbit anti-pPDH (37115, CST), rabbit anti-Glyt2 (MA552662, Invitrogen), mouse anti-GAD67/GAD1 (ab26116, Abcam), rabbit anti-CaMKII (ET1608-47, HUABIO). Sections were then washed three times in PBS for 5 minutes each and incubated with fluorescent-conjugated secondary antibodies for 1 hour at room temperature. The secondary antibodies that were used are Alexa Fluor 488-conjugated donkey anti-rabbit IgG (20015, Biotium), Alexa Fluor 568-conjugated donkey anti-mouse IgG (20105, Biotium), and Alexa Fluor 633-conjugated donkey anti-mouse IgG (20124, Biotium). After washing three times by PBS (5 minutes each), the sections were counterstained with DAPI (ZLI-9557, ZSGBBIO). Images were captured by either 20×magnification using a slide scanner (TissueFAXS, TISSUEGNOSTICS) or captured by 40× magnification using FV1000 laser confocal microscope (Olympus). Following IF, positive cells for c-Fos, and positive area for pPDH were counted using the TissueQuest (TISSUEGNOSTICE) (*62*).

**Western blot analysis**

Western blot analysis was performed to assess NR1 protein levels in the anterior and posterior cerebellar lobules. Total protein was extracted from mouse brain tissues using a lysis buffer (89900, Sigma), and concentrations were determined via a bicinchoninic acid (BCA) assay (23227, Sigma). Equal amounts of protein were separated by sodium dodecyl sulfate–polyacrylamide gel electrophoresis (SDS-PAGE) and transferred onto polyvinylidene fluoride (PVDF) membranes (Yangguang Bio).

Following transfer, membranes were blocked with 5% bovine serum albumin (BSA) and incubated overnight at 4 °C with the following primary antibodies: mouse anti-GAPDH (ab8245, Abcam) and mouse anti-NR1 (05-432, Sigma). The membranes were subsequently incubated for 1.5 h at room temperature (20–24 °C) with a horseradish peroxidase (HRP)-conjugated goat anti-mouse IgG secondary antibody (ZB-2305, ZSGB-BIO). Protein bands were visualized using a chemiluminescent imaging system (SAGECREATION), and signal intensities were quantified using ImageJ 2.0.0 software (NIH, Bethesda, MD, USA). NR1 expression levels were normalized to GAPDH as an internal loading control.

**Stereotaxic injection**

Mice were anesthetized using 1% pentobarbital (50 mg/kg) during surgical procedures. Hair was subsequently removed, the skin was cleaned, and the skull was dried. Viral injections were performed at a rate of 50 nL/min, with injection needles remaining in place for an additional 10 minutes to facilitate diffusion. To target different cerebellar lobes, using the following stereotaxic coordinates: lobule III (AP: -5.38 mm, ML: 0.0 mm, DV: -1.25 mm), lobule IV/V (AP: -6.36 mm, ML: 0.0 mm, DV: -1.27 mm), lobule VI (AP: -6.84 mm, ML: 0.0 mm, DV: -1.00 mm), lobule VII (AP: -6.84 mm, ML: -2.5 mm, DV: -1.65 mm), lobule VIII (AP: -6.96 mm, ML: 0.0 mm, DV: -1.65 mm), and lobule IX (AP: -7.08 mm, ML: 0.0 mm, DV: -2.0 mm). Other site coordinates include: LaVe (AP: -6.48 mm, ML: 1.5 mm, DV: -2.9 mm), Rt (AP: -7.56 mm, ML: 1.25 mm, DV: -4.65 mm), Pn (AP: -4.48 mm, ML: 0.5 mm, DV: -4.65 mm).

**Fiber photometry**

Fiber photometry is a crucial technique for investigating brain-behavior relationships in vivo, as established in prior studies *(63)*. To record calcium signal changes in cerebellar GCs using optical fiber photometry in the 5-CSRTT model, *Atoh1-Cre* mice were injected with 400 nL of AAV-hSyn-DIO-GCaMP6s (PT-0091, BrainVTA) and into one of several cerebellar lobes, ranging from lobule III to lobule IX. Fiber-optic inserts (200 μm, NA = 0.37; Originopto) were then implanted 0.02 mm above the viral injection sites. These fibers were secured to the skull using screws and dental cement, as previously described (*41*).

Fiber photometry data acquisition was carried out using a Neurophotometrics fiber photometry system (Biolinkoptics). This system utilized a continuously illuminated 470-nm blue-light LED with an intensity range of 40-100 μW to excite the GCaMP. A dichroic mirror was incorporated into the fluorescence light path to direct the emitted green fluorescence toward a photomultiplier tube (PMT, R3896). The fluorescence signals captured by the PMT were processed using MATLAB (Mathworks).

Fluorescence change (ΔF/F) values were derived by calculating (F−F0)/F0, where F0 represents the average baseline fluorescence signal recorded before the test. ΔF/F values were presented as heatmaps or average plots, with the shaded area representing the standard error of the mean (SEM). To assess the relationship between neural activity and physical behavior, simultaneous recordings of calcium signals and video tracking were obtained. Photometry signals were aligned to specific behavioral events of interest in the 5-CSRTT, including correct response, incorrect response, omission, and premature response, to perform event-aligned analysis. Subsequently, summary statistics were calculated by measuring the area under the curve (AUC) for each event type within a defined time interval (ranging from -5 s to 10 s relative to the event onset).

**Chemogenetics manipulation**

For chemogenetic manipulation of GCs in lobules IV/V or VI, *Atoh1-Cre* mice were injected with 400 nL of AAV-EF1α-DIO-hM3D(Gq)-EGFP (BC-0144, BrainVTA), AAV-EF1α-DIO-hM4D(Gi)-EGFP (BC-0623, BrainVTA), or AAV-EF1α-DIO-EGFP (control, BC-0015, Braincase). To achieve pathway-specific chemogenetic targeting of cerebellar-afferent projections, a dual-virus strategy was employed. First, C57BL/6J mice received a 400 nL injection of AAV-retro-Cre-mCherry (BC-0539, Braincase) into the granular layer of either lobule VI or IV/V. Subsequently, Cre-dependent AAVs (300 nL of AAV-DIO- hM4Di or AAV-DIO-hM3Dq) were injected into the corresponding upstream nuclei, namely the Pn, LaVe, or Rt. Furthermore, to specifically activate the Rt-lobule IV or Pn-lobule VI circuits, *DAT-HET* mice were injected with 400 nL of AAV-retro-Cre-mCherry into the granular layer of lobule VI or IV/V, followed by the delivery of AAV-DIO-hM3Dq or AAV-DIO-EGFP into the Rt or Pn. Behavioral testing commenced approximately 45 days post-surgery, once mice achieved consistent baseline performance levels. Task performance was assessed 30 minutes after intraperitoneally (*i.p.*) injected with either the selective DREADD ligand clozapine-N-oxide (CNO) (C0832, Sigma) at a dose of 3 mg/kg or 0.9% saline solution. Validation of the virus injection site and treated as described above.

**Spatial library preparation and sequencing**

*Spatial library preparation and sequencing*

Tissue sectioning, Topuidine Blue staining, imaging, and first permeabilization were consistent with the user guide of the BMKMANU S1000 Tissue Optimization Kit (ST03003, BMKMANU). The library was constructed according to the BMKMANU S1000 Library Construction Kit. Library was checked on Qubit to conduct quality control. The library was sequenced with Novaseq 6000 using PE150 with an average of 876 million reads per sample. The raw sequencing data were mapped to the reference genome (mm10) by BSTMatrix (version 2.0) using default parameters. The expression matrix was transformed into a Seurat object using Seurat package (version 4.0.1) (*64*). The NormalizeData function was used to normalize the gene counts across all samples. We ﬁrstly identified the top 2,000 genes with the highest variability by using the FindVariableGenes function. To eliminate batch effects, we used the FindIntegrationAnchors and IntegrateData functions to integrate Naive and 5-CSRTT samples using all detected genes. Subsequently, we reduced the dimensionality of our data through principal component analysis (PCA) and t-distributed Stochastic Neighbor Embedding (t-SNE) using RunPCA and the RunTSNE functions, respectively. Clustering was conducted using FindClusters function with default parameters, and resolution parameters set from 0.5 to 2.5. To pinpoint marker genes for each cluster, we utilized the FindMarkers function. To annotate cell clusters to known cell types, we evaluated the expression patterns of cell type-specific marker genes that identified from previous literature (*65, 66*)

*Rank-rank hypergeometric overlap analysis*

Rank-rank hypergeometric overlap (RRHO) is a threshold-free method designed to identify and visually represent the overlap of genes that are altered in the same or opposite directions across two datasets (*67*). We first calculate the log2 fold change values of genes between pairs of brain regions. Those exhibiting upregulation were positioned at the top of the list, downregulated genes at the bottom, and genes with no change were found in the middle. Utilizing a heatmap generated through the odds ratio approach, we can effectively illustrate the intensity, distribution, and correlation thresholds that define the relationships between different brain regions. RRHO analysis was performed using RRHO package (version 1.26.0).

*Gene functional enrichment analysis*

Differentially expressed genes (DEGs, two-sided Wilcoxon-test, log fold-change threshold = 0.25, FDR < 0.05) of each brain region were identified and subsequently used for Gene Ontology (GO) term enrichment analysis using clusterProfiler package (version 3.18.1) (*68*). The enriched GO terms of DEGs were selected and visualized utilizing ggplot2 package (version 3.4.2). To predict the biological relationship of DEGs of GCs in AL or PL regions, Cytoscape (version 3.7.1) (*69*) plug-in ClueGO (version 2.5.9) (*70*) was used. Only terms at a significant level Bonferroni adjusted p value ≤ 0.05 were kept. We used Tau variation to evaluate the specificity of DEGs. We evaluated CV variation to evaluate the variability of DEGs.

The Tau variation is defined as:

$$Tau=\frac{\sum_{i=1}^{n} (1-\frac{E_{i}}{\max_{E\in S} f(E)})}{n-1}$$

Where *n* is the number of cells and *E*_i_ is the expression value of gene in cell *i*. *S* is the entire dataset of expression values in all cells.

$$CV=\frac{\sqrt{\frac{1}{n-1}\sum_{i=1}^{n} {(E_{i}-\overline{E})}^{2}}}{\overline{E}}$$

Where *n* is the number of cells, *E*_i_ is the expression value of gene in cell *i* and $\overline{E}$ is the average expression level.

*Spatial cellular communication analysis*

We used both CellChat (*71*) and dsCellNet (*72*) to infer cellular communication among different cell types in Naïve and 5-CSRTT, respectively (adjusted p-value < 0.05). The importance score of each cell type is calculated as the scaled sum of outgoing interactions using dsCellNet. To extract the 5-CSRTT associated cellular communication, we subtracted the interactions of 5-CSRTT from the interactions of Naïve. The subtracted cellular communication network was visualized by Cytoscape (version 3.7.1).

**RNAscope *in situ* hybridization**

Mice were anesthetized using 1% pentobarbital (50 mg/kg) and subsequently perfused with saline maintained at 4°C. The brain was immediately dissected on ice, embedded in an optimal cutting temperature (OCT) medium, rapidly frozen, and stored at -80°C. Sagittal sections of the cerebellar vermis were cut at a 10 μm thickness using a cryotome (Thermo Fisher Scientific). The RNAscope in situ hybridization (ISH) was performed using the Multiple Fluorescent Detection Reagents v2 kit (323110, ACDbio). Sections without OCT embedding were baked at 60°C for 1 hour, followed by fixation in 10% neutral buffered formalin for 40 minutes. The fixed sections were dehydrated in sequential ethanol concentrations (50%, 70%, and 100%) and treated with H_2_O_2_ for 10 minutes, followed by protease III treatment for 15 minutes. Subsequent steps followed the manufacturer’s protocol (*73*). For all experiments, Probe-Mm-*Grin1* (431611, ACDbio) and Opal 520 reagent (OP-001001, Akoya Biosciences) were used to detect *Grin1* in the green fluorescence channel. Images post-RNAscope ISH were captured using a TissueFAXS imaging system (TISSUEGNOSTICS) with a 40× objective lens, maintaining consistent imaging parameters as defined by both positive and negative controls (using Positive Control Probe-Mm, 320881, and Negative Control Probe-Mm, 320871, ACDbio).

For quantitative analysis, the ratio of “*Grin1* mRNA / DAPI” was calculated by counting the *Grin1* mRNA puncta along with DAPI-stained puncta within different granule cell layers using the IF2 APP in StrataQuest (TISSUEGNOSTICS). An independent, experienced researcher, blinded to study, performed all the counts.

**Cannula infusion experiment**

A 26-gauge single guide cannula (O.D. 0.48 mm) (RWD Life Science) was inserted into the cerebellar granule cell layer of lobule Ⅳ/Ⅴ or lobule Ⅵ. A compatible 26-gauge single cap (O.D. 0.30 mm) was inserted into the guide cannula to prevent clogging during the recovery period. The cannulas were secured to the skull by using stainless steel screws, and the assembly was fixed in dental cement. After mice had recovered for at least 7 days, drugs were microinjection with a 30-gauge single injector cannula. The procedure follows previously established protocols (*74*).

The noncompetitive NMDA receptor antagonist dizocilpine (20 or 40 nM, 1 μL; MK-801, Sigma) (*24*) or NMDA (50 μM, 1 μL; HY-17551, MCE) (*75*) was dissolved in sterile 0.9% saline before the experiments. Before the local drug infusion, 30-gauge single injector cannula were inserted into the guide cannula to ensure clear passage and then pull out. One microlitre of drug was infused (0.2 μL/min) through another set of 25-gauge single plug, which were connected to the microsyringe (CHEMYX) with Polyethylene pipe. After taken enough drug, the injector cannula was kept in place for an additional 5 min to allow an adequate local drug diffusion and minimize spread of the drug along the injection track. The 5-CSRTT or OFT was performed 30 min after the injection of MK801 or NMDA. To check the drug infusion sites, mice were injected with 1 μL Dil (5 μM) (C1036, Beyotime) into each cerebellar lobules after all behavioral procedures. Fluorescent image acquisition was performed with the TissueFAXS slide scanning system (TISSUEGNOSTICS). Only data from mice with correctly sited injections were used.

**Viral tracing**

*Tracing the upstream connections of cerebellar GCs*

*Atoh1-Cre* mice were injected with 100 nL of a mixture of AAV vectors (1:1, AAV2/8-EF1α-DIO-△RVG; AAV2/8-EF1α-DIO-EGFP-T2A-TVA) (PT-0023; PT-0062, Brain VTA) into lobule IV/V or VI of cerebellum. Three weeks post-AAV injection, the mice were further injected at the same cerebellar sites with RV-EnvA-△G- mCherry (BC-RV-EnvA-844, Brain VTA). One week after the RV injection, the mice were sacrificed, and their brains were processed for histological analysis. To validate the virus injection site and quantify the rabies tracing results, the brains were isolated 7 days after RV injection and processed as described in previously published protocols (*25*). Quantification of subregions relied on boundaries outlined in a mouse brain atlas. Anatomically contiguous regions with similar functions and lacking clear-cut anatomical boundaries were combined (*76*). For example, the pontine nuclei (Pn), including its caudal (PnC), oral (PnO), and ventral (PnV) parts were combined into Pn; the cerebral peduncle (cp), including its inferior (icp) and middle (mcp) parts were combined into cp; The gigantocellular nucleus (Gi), including its alpha (GiA) and ventral (GiV) parts, and the lateral (LPGi) and dorsal (DPGi) paragigantocellular nuclei were combined into GRt (gigantocellular reticular nucleus); The reticular nucleus (Rt) including its intermediate (IRt) and lateral (LRt), as well as its rostroventrolateral (RVL) parts; The parvicellular reticular nucleus (PCRt) and its alpha part (PCRtA) were combined into PRt (parvicellular reticular nucleus); The medullary reticular nucleus (MRt) including its dorsal (MdD) and ventral (MdV) parts; The paramedian reticular nucleus (PMn) and paramedian raphe nucleus (PMnR) were combined into PMn; The cuneate nucleus (Cu) and its external part were combined into Cu; The Principal sensory trigeminal nucleus (Pr5) and its sorsomedial (Pr5DM) and ventrolateral (Pr5VL) parts were combined into Pr5; and the oral (Sp5O), dorsomedial (DMSp5), interpolar (Sp5I), and caudal (Sp5C) parts of the spinal trigeminal nucleus (Sp5) were combined into Sp5; The medial vestibular nucleus (MVe), including its magnocellular (MVeMC) and parvicellular (MVePC) parts, as well as the vestibulocerebellar nucleus (VeCb) were combined into MeVe (medial vestibular nucleus); The lateral (LVe), spinal (SpVe), and superior (SuVe) of vestibular subnuclei, and nucleus X (X) were combined into LaVe (lateral vestibular nucleus); The alpha (SubCA), dorsal (SubCD) and ventral (SubCV) parts of subcoeruleus nucleus were combined into SubC (subcoeruleus nucleus). Brain subregions in which no more than two mice found projection cells in the same subregion were not included in the final analysis.

*Tracing the downstream connections of LaVe, Rt or Pn*

C57BL/6J mice were injected with 80 nL AAV2/1-hSyn-H2B-mclover3-pA (S0864, Taitool) into LaVe, Rt or Pn. Four weeks post-AAV injection, the mice were sacrificed, and their brains were processed for histological analysis.

*Tracing the upstream connections of Rt or Pn*

C57BL/6J mice were injected with 100 nL AAV/Retro-mCherry (BC-0539, Braincase) or AAV/Retro-EGFP (PT-0156, BrainVTA) into Rt or Pn. Three weeks post-AAV injection, the mice were sacrificed, and their brains were processed for histological analysis. Quantification of subregions relied on boundaries outlined in a mouse brain atlas.

**Quantification and Statistical Analysis**

Statistical analyses were performed using GraphPad Prism 10.1.2. Data were presented as mean ± SEM. The accepted level of significance was p < 0.05. Two-tailed Student’s t-test, repeated measure, One-way or Two-way ANOVA were used for statistical analysis.

Supplementary Text

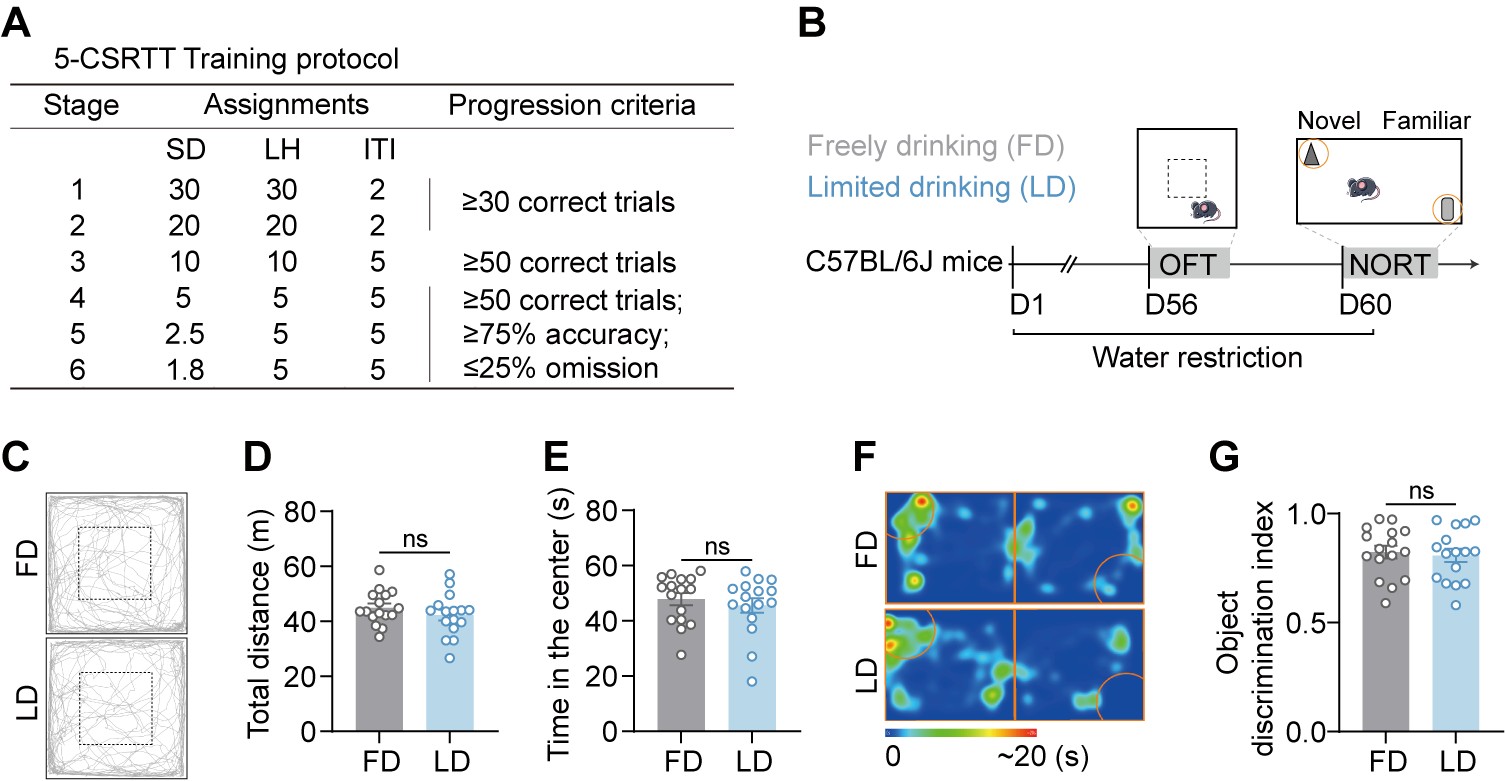

Fig. S1. 5-CSRTT training protocol and water restriction controls, related to Fig. 1.

(A) Parameters and progression criteria for the six-stage 5-CSRTT training protocol. SD, stimulus duration; LH, limited hold; ITI, inter-trial interval.

(B) Experimental timeline for assessing the potential side effects of the water restriction regimen (n = 16). Mice were divided into freely drinking (FD) and limited drinking (LD) groups before undergoing behavioral assays.

(C–E) Water restriction does not alter locomotor activity or anxiety-like behavior. (C) Representative track plot in the open field test (OFT). (D and E) Quantification of total distance traveled (D), and time spent in the center area (E).

(F and G) Water restriction does not impair recognition memory. (F) Representative heatmaps of exploration trajectories in the novel object recognition test (NORT) for FD (top) and LD (bottom) groups. (G) Quantification of the object discrimination index.

Data are presented as mean ± SEM. ns, not significant by unpaired t-test.

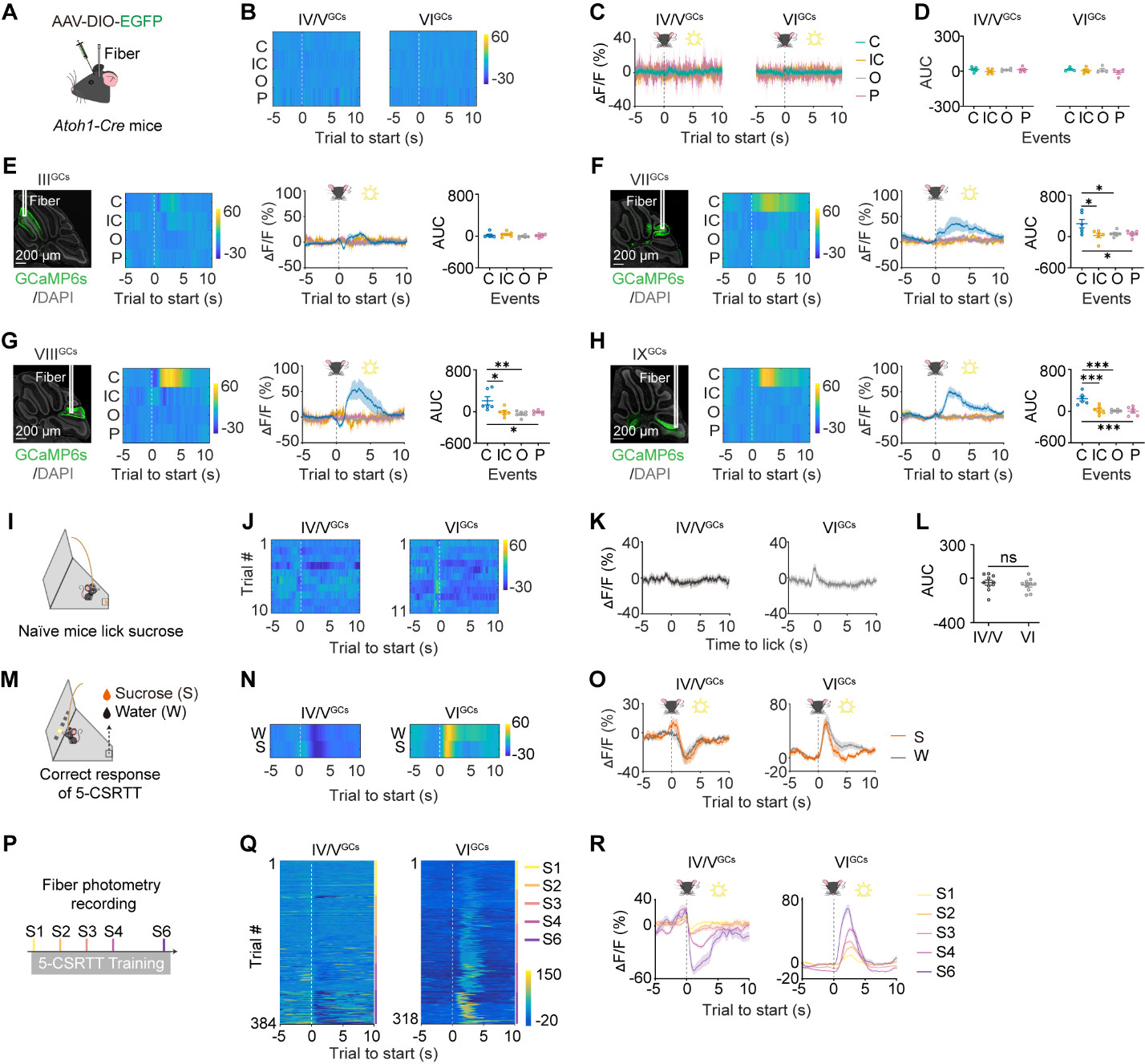

Fig. S2. Lobule-specific calcium dynamics in granule cells during attention, reward, and learning, related to Fig. 1.

(A–D) EGFP controls show no motion artifacts. (A) Diagram of *Atoh1-Cre* mice with AAV-DIO-EGFP. (B) Heatmaps of fluorescence changes in EGFP-expressing mice during 5-CSRTT. (C) Average ΔF/F traces showing stable fluorescence across all outcomes. (D) Quantification of AUC confirms no significant signal changes in EGFP controls. n = 4-5 *Atoh1-Cre* mice.

(E–H) Ca^2+^ activity in other cerebellar lobules during the 5-CSRTT. Representative images showing GCaMP6s expression and fiber placement, heatmaps of average Ca^2+^ signals (ΔF/F) aligned to trial start and average ΔF/F traces for correct (C), incorrect (IC), omission (O), and premature (P) trials, and quantification of AUC (-5 to 10 s) in lobule III (E), VII (F), VIII (G), and IX(H) GCs. Shading indicates SEM. n = 6 *Atoh1-Cre* mice.

(I–L) GCs show minimal response to free reward consumption. (I) Schematic of fiber photometry recording in naïve mice licking sucrose. (J) Heatmaps of Ca^2+^ activity in lobule IV/V and VI GCs aligned to first lick. (K) Average ΔF/F traces during sucrose consumption. (L) Quantification of AUC shows no significant activation or suppression during free reward consumption. n = 10 *Atoh1-Cre* mice.

(M–O) Attentional signals are robust to reward devaluation. (M) Schematic of the reward devaluation experiment (sucrose vs. water) in the 5-CSRTT. (N) Heatmaps of Ca^2+^ activity during correct trials rewarded with sucrose (S) or water (W). (O) Average ΔF/F traces showing that the characteristic suppression (IV/V) and activation (VI) patterns persist regardless of reward type.

(P–R) Attentional signals are learning-dependent. (P) Schematic of longitudinal fiber photometry recording across training stages (S1–S6). (Q) Heatmaps of Ca^2+^ activity in lobule IV/V and VI GCs during correct trials across training stages. Note the emergence of the suppression (IV/V) and activation (VI) patterns only in later stages. (R) Average ΔF/F traces at different training stages. Pre-training (Stage 1) shows minimal modulation compared to Post-training (Stage 6). Data are presented as mean ± SEM. **P* < 0.05, ***P* < 0.01, ****P* < 0.001; ns, not significant by One-way ANOVA with Tukey's multiple comparisons test (D-H), or unpaired t-test (L).

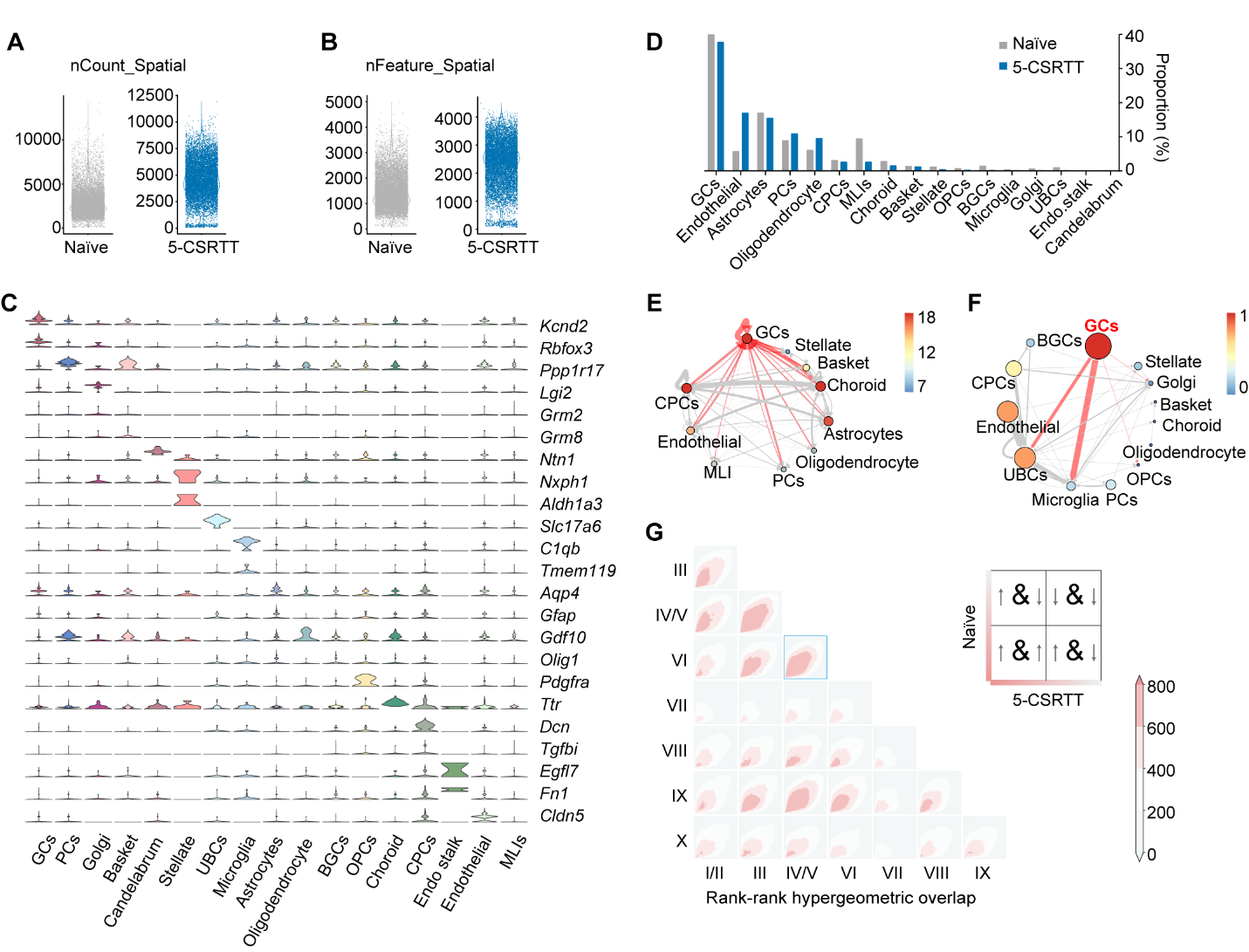

Fig. S3. Quality control, cell type annotation, and cellular interaction analysis of spatial transcriptomics data, related to Fig. 3

(A and B) Quality control metrics for spatial transcriptomic sequencing. Violin plots showing the distribution of total unique molecular identifier (UMI) counts (A, nCount_Spatial) and detected gene numbers (B, nFeature_Spatial) per spot in Naïve and 5-CSRTT samples.

(C) Validation of cell type identity. Stacked violin plots displaying the expression levels of canonical marker genes used to annotate the 17 identified cell clusters. Key markers include *Kcn2* and *Rbfox3* for GCs, *Itpr1* for PCs, and *Gfap* for astrocytes. Detail see Table S2.

(D) Cellular composition of the cerebellar vermis. Bar graph showing the proportion of each cell type in Naïve (grey) and 5-CSRTT (blue) mice. GCs constitute the most abundant population (~40%).

(E and F) GCs are central hubs in the attention-modulated communication network. (E) Ligand-receptor interaction network inferred by CellChat. The network depicts interactions specific to the 5-CSRTT condition (subtracted by Naïve baseline). Node size represents interaction number; edge width represents interaction strength. GCs (red node) exhibit the highest connectivity. (F) Ligand-receptor interaction network inferred by dsCellNet, confirming the central role of GCs in the cellular crosstalk associated with the 5-CSRTT.

(G) Transcriptomic concordance across lobules. Rank-rank hypergeometric overlap (RRHO) heatmaps comparing the gene expression signatures of GCs across different cerebellar lobules. The color intensity represents the degree of overlap in gene ranking. The highlighted box (blue outline) indicates the specific transcriptomic relationship between Lobule IV/V and Lobule VI.

Data are presented as mean or representative plots.

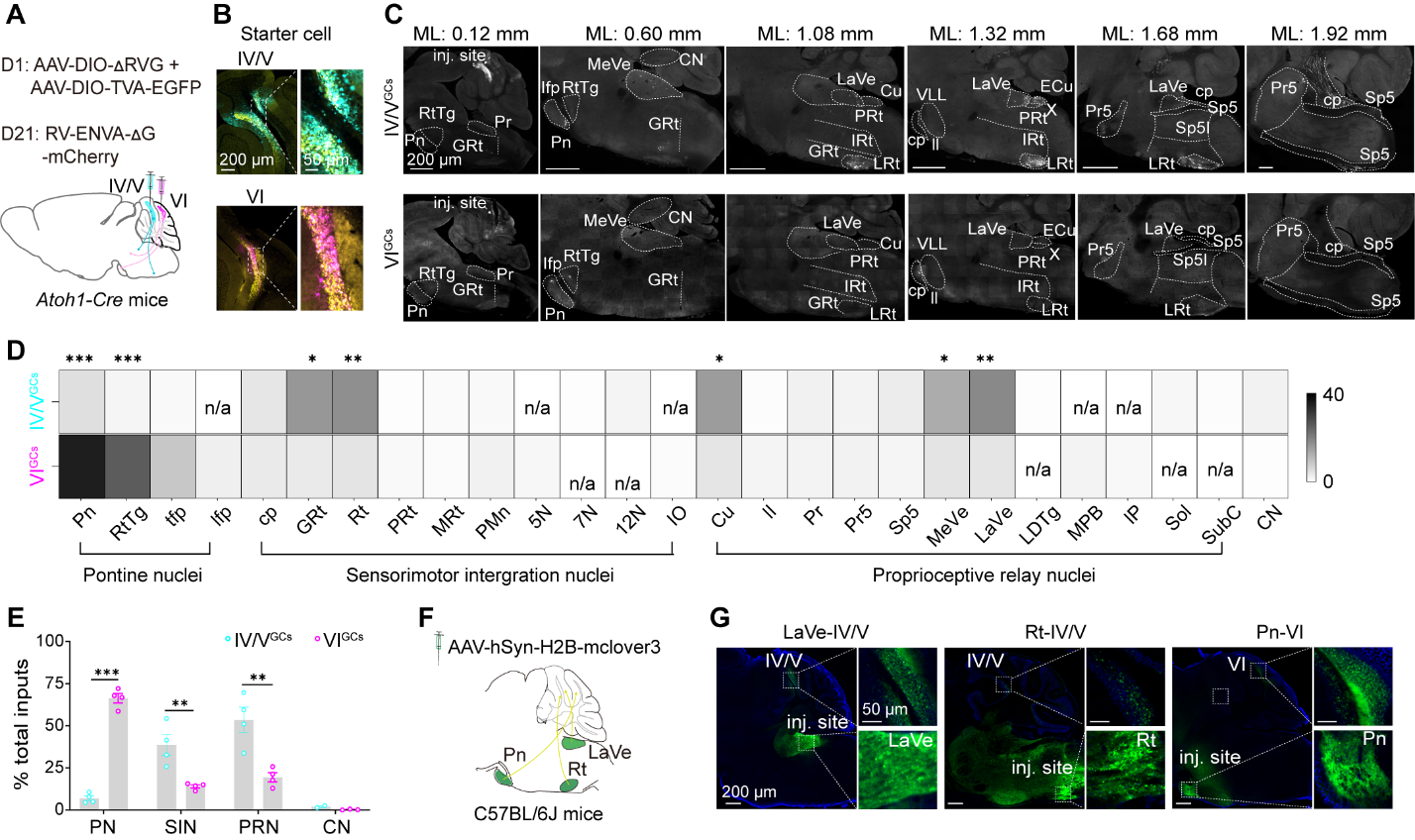

Fig. S4. Anatomical segregation of upstream inputs targeting anterior and posterior cerebellar GCs, related to Fig. 5.

(A) Schematic of the monosynaptic retrograde tracing strategy. *Atoh1-Cre* mice were injected with helper viruses (AAV-DIO-TVA-EGFP and AAV-DIO-ΔRVG) followed by pseudotyped rabies virus (RV-ENVA-ΔG-mCherry) into lobule IV/V or VI to map inputs to GCs. n = 4, *Atoh1-Cre* mice.

(B) Representative confocal images of the injection sites showing starter cells co-expressing EGFP (yellow) and mCherry (cyan or purple) in lobule IV/V (top) and VI (bottom). Scale bars, 200 μm (low mag), 50 μm (high mag).

(C) Representative sagittal brain sections showing the distribution of monosynaptic input neurons (mCherry^+^) across the brainstem. Note the distinct innervation patterns: Lobule IV/V GCs receive dense inputs from the vestibular (LaVe, MeVe) and reticular (Rt, GRt) nuclei, whereas Lobule VI GCs are primarily innervated by the pontine nuclei (Pn). Scale bars, 200 μm.

(D) Heatmap profiling the specific contribution of individual brainstem nuclei to the total input pool for lobule IV/V (top row) versus lobule VI (bottom row). Darker shades indicate higher percentage of input.

(E) Quantification of total input fractions categorized into Pontine, Sensorimotor intergration (SIN), Proprioceptive relay (PRN), and Cerebellar Nuclei (CN) groups. Lobule VI GCs are dominated by pontine inputs, while Lobule IV/V GCs receive predominantly motor and sensory inputs.

(F) Schematic of the anterograde tracing strategy for validation. AAV-hSyn-H2B-mclover3 was injected into the Lateral Vestibular Nucleus (LaVe), Reticular Nucleus (Rt), or Pontine Nuclei (Pn).

(G) Representative images of EGFP^+^ axonal terminals in the cerebellar cortex. Axons from LaVe and Rt preferentially terminate in the granular layer of lobule IV/V, whereas axons from Pn preferentially target lobule VI. Scale bars, 200 μm (low mag), 50 μm (high mag).

Abbreviations: Pn, pontine nuclei; RtTg, reticulotegmental nucleus of the pons; tfp, transverse fibers of the pons; lfp, longitudinal fasciculus of the pons; cp, cerebral peduncle; GRt, gigantocellular reticular nucleus; Rt, reticular nucleus; PRT, parvicellular reticularnucleus; MRt, medullary reticular nucleus; PMn, paramedian reticular nucleus; 5N, motor trigeminal nucleus; 7N, facial nucleus; 12N, hypoglossal nucleus; IO, inferior olivary nucleus; Cu, cuneate nucleus; ll, lateral lemniscus; Pr, prepositus nucleus; Pr5, principal sensory trigeminal nucleus; Sp5, spinal trigeminal tract; MeVe, medial vestibular nuclei; LaVe, vestibular nucleus; LDTg, laterodorsal tegmental nucleus; MPB, medial parabrachial nucleus; IP, interpeduncular nucleus; Sol, solitary nucleus; SubC, subcoeruleus nucleus.

Data are presented as mean ± SEM (n = 4 mice). **P* < 0.05, ***P* < 0.01, ****P* < 0.001 by unpaired t-test (D) or Two-way ANOVA with Sidak's multiple comparisons test (E).

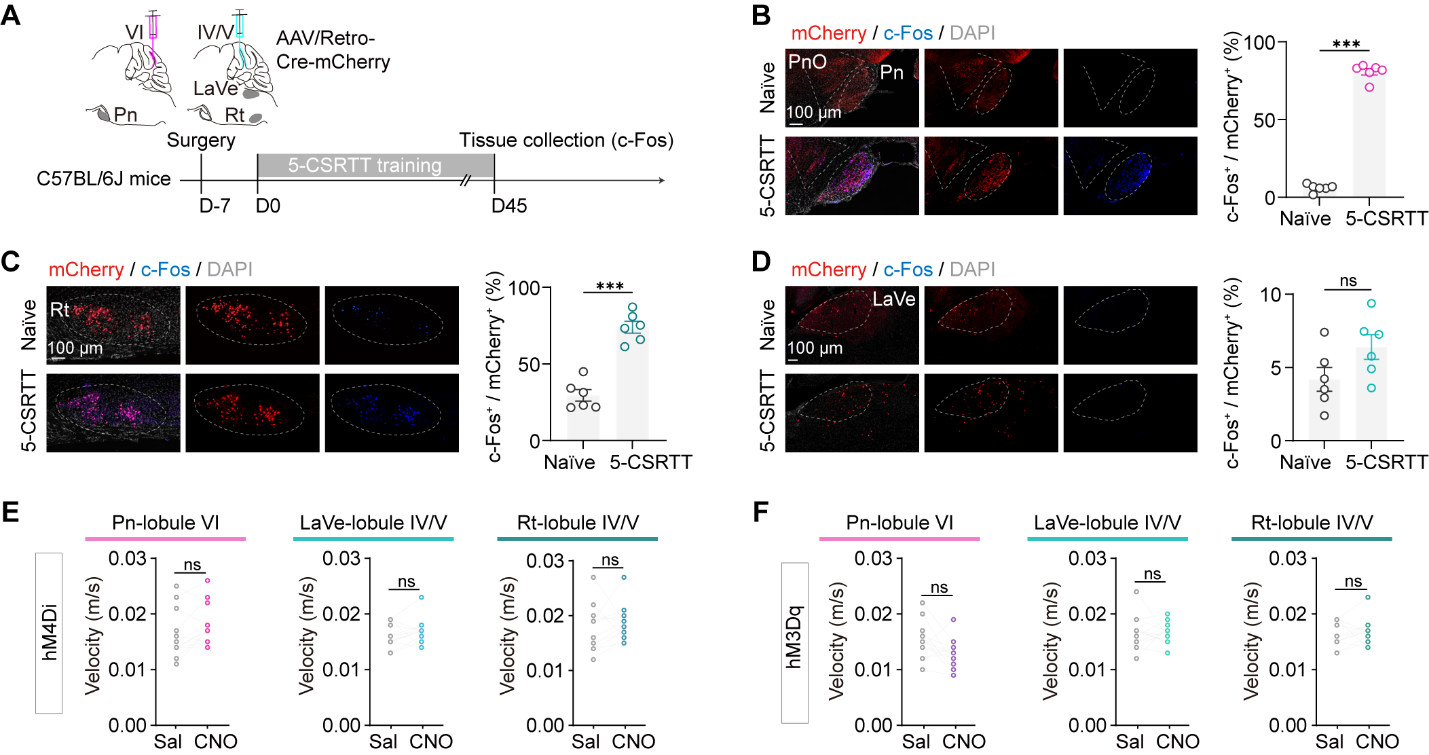

Fig. S5. Task-dependent activation of cerebellar afferents and motor control for circuitry manipulations, related to Fig. 5.

(A) Experimental design for mapping the activity of upstream projection neurons. AAV-Retro-Cre-mCherry (red) was injected into cerebellar lobule VI or IV/V to retrogradely label projection neurons in the Pontine Nuclei (Pn), Lateral Vestibular Nucleus (LaVe), or Reticular Nucleus (Rt). c-Fos expression was assessed in Naïve versus 5-CSRTT trained mice.

(B–D) Selective recruitment of specific cerebellar afferents during attention. (B) Representative confocal images and quantification of c-Fos expression (blue) in Pn neurons projecting to lobule VI (mCherry^+^). Scale bar, 100 μm. (C) Analysis of Rt neurons projecting to lobule IV/V. (D) Analysis of LaVe neurons projecting to lobule IV/V.

(E–F) Pathway-specific chemogenetic manipulation does not alter locomotor speed. Average velocity during the 5-CSRTT for mice expressing hM4Di (E) or hM3Dq (F) in the Pn → Lobule VI (left), LaVe→ Lobule IV/V (middle), or Rt→ Lobule IV/V (right) pathways.

Data are presented as mean ± SEM. n = 6 mice for c-Fos (B–D); n = 12 mice for behavior (E). **P* < 0.05, ***P* < 0.01, ****P* < 0.001; ns, not significant (Unpaired t-test for B–D; Paired t-test for E–F).

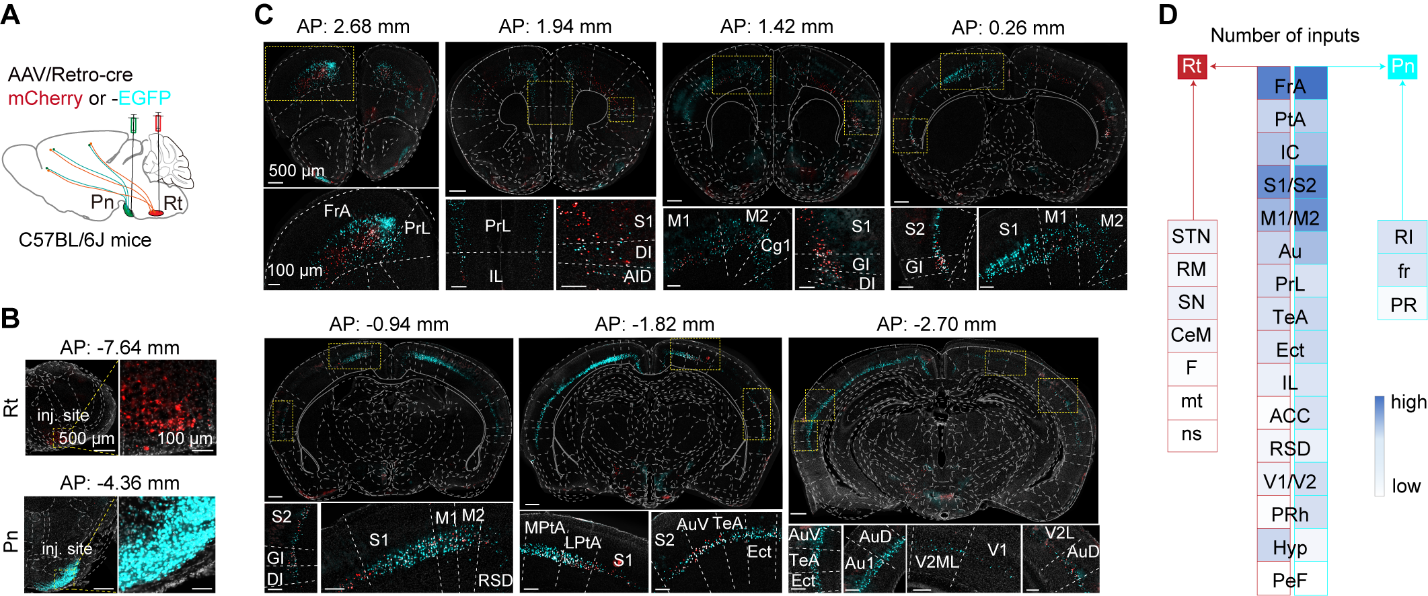

Fig. S6. Convergent cortical projections to the Pn and Rt.

(A) Schematic illustration of the dual retrograde tracing strategy used to map and compare upstream inputs. AAV-Retro-EGFP (cyan) and AAV-Retro-mCherry (red) were injected into the Pontine Nuclei (Pn) and Reticular Nucleus (Rt), respectively.

(B) Representative confocal images showing the injection sites and viral expression of the starter cells in the Pn (bottom, cyan) and Rt (up, red). Scale bars, 500 μm (left), 100 μm (right).

(C) Representative coronal brain sections displaying the anatomical distribution of retrogradely labeled neurons. Images depict prominent labeling of both Pn-projecting (cyan) and Rt-projecting (red) neurons within overlapping regions of the frontal cortex (FrA, PrL, M2) and parietal/sensorimotor cortex (M1, S1, Cg1). Scale bars, 500 μm (low mag), 100 μm (high mag).

(D) Schematic summary and heatmap quantification of input densities from various brain regions. The heatmap illustrates the relative number of input neurons targeting the Pn versus the Rt, highlighting shared innervation from frontal and parietal cortical areas (FrA, M1/M2, S1/S2) alongside distinct subcortical contributions.

Data are presented as representative images and quantification from n = 3 mice.

**Abbreviations:** FrA, frontal association cortex; PtA, parietal association cortex; IC, insular cortex; S1/S2, primary/secondary somatosensory cortex; M1/M2, primary/secondary motor cortex; Au, auditory cortex; PrL, prelimbic cortex; TeA, temporal association cortex; Ect, ectorhinal cortex; IL, infralimbic cortex; ACC, anterior cingulate cortex; RSD, retrosplenial dysgranular cortex; V1/V2, primary/secondary visual cortex; PRh, perirhinal cortex, Hyp, hypothalamic nucleus; PeF, perifornical nucleus; STN, subthalamic nucleus; RM, retromammillary nucleus; SN, substantia nigra; CeM, central amygdaloid nucleus, medial division; F, nucleus of the fields of Forel; mt, mammillothalamic tract; ns, nigrostriatal tract; RI, rostral interstitial nucleus; fr, fasciculus retroflexus; PR, prerubral field.

**Table S1. Primers used for genotyping**

| **Mouse line** | **Primer sequence (5’-3’)** |
| --- | --- |
| *Atoh1-Cre* | CCGGCAGAGTTTACAGAAGC |
|  | CCGGCAGAGTTTACAGAAGC |
| *DAT-HET* | GTTGATGAGGGTGGAGTTGGTC |
|  | GCCGCATAACCAGTGAAACAGC |
|  | TCCATAGCCAATCTCTCCAGTC |

Table S2. Annotated gene marker for cell types in spatial transcriptomic sequencing, related to Fig. 3B

| **Cell type** | **Gene marker** | |
| --- | --- | --- |
| GCs | *Kcnd2* | *Rbfox3* |
| Purkinje cell (PCs) | *Ppp1r17* |  |
| Golgi cell | *Lgi2* | *Grm2* |
| Basket cell | *Grm8* |  |
| Stellate cell | *Ntn1* |  |
| Candelabrum cell | *Nxph1* | *Aldh1a3* |
| Unipolar brush cell (UBCs) | *slc17a6* |  |
| Microglia | *C1qb* | *Tmem119* |
| Astrocytes | *Aqp4* | *Gfap* |
| Bergmann glia cell (BGCs) | *Gdf10* |  |
| Oligodendrocyte | *Olig1* |  |
| Oligodendrocyte precursor cell (OPCs) | *Pdgfra* |  |
| Choroid | *Ttr* |  |
| Choroid plexus cell (CPCs) | *Dcn* | *Tgfbi* |
| Endo.stalk | *Egfl7* | *Fn1* |
| Endothelial | *Cldn5* |  |
| MLIs (molecular layer interneuron) | unknown |  |

Movie S1.

Demonstration of the touch-screen version of the 5-choice serial reaction time task (5-CSRTT).

Movie S2.

Calcium signal dynamics of granule cells (GCs) in the anterior cerebellar vermis (Lobules IV/V) during 5-CSRTT performance.

Movie S3.

Calcium signal dynamics of granule cells (GCs) in the posterior cerebellar vermis (Lobule VI) during 5-CSRTT performance.

Data S1.

Tabulated raw data and statistical analyses underlying all figures.
